# Neural temporal context reinstatement of event structure during memory recall

**DOI:** 10.1101/2021.07.30.454370

**Authors:** Lynn J. Lohnas, M. Karl Healey, Lila Davachi

## Abstract

Although life unfolds continuously, experiences are generally perceived and remembered as discrete events. Accumulating evidence suggests that event boundaries disrupt temporal representations and weaken memory associations. However, less is known about the consequences of event boundaries on temporal representations during retrieval, especially when temporal information is not tested explicitly. Using a neural measure of temporal context extracted from scalp electroencephalography, we found reduced temporal context similarity between studied items separated by an event boundary when compared to items from the same event. Further, while participants free recalled list items, neural activity reflected reinstatement of temporal context representations from study, including temporal disruption. A computational model of episodic memory, the Context Maintenance and Retrieval model (CMR; Polyn, Norman & Kahana, 2009), predicted these results, and made novel predictions regarding the influence of temporal disruption on recall order. These findings implicate the impact of event structure on memory organization via temporal representations.

## Introduction

Building on centuries of philosophy and psychology, an overarching question in cognitive neuroscience concerns the transformation of the objective, external environment into subjective, internal representations. This question is central to characterizing how memory representations are stored, reinstated and organized, which in turn influences how memory biases ongoing cognition. The relationship between temporal perception and memory organization is a ubiquitous example of this point. Time moves forward constantly and continuously, yet temporal information is rarely perceived or remembered in this way. When reflecting on one’s day, it is more likely that information will be remembered as subjective but meaningful and distinct events (e.g., ‘I prepared breakfast’, ‘I ate breakfast’), rather than by objective time only (e.g. ‘From 8:00-8:15am I prepared breakfast and started eating breakfast’, ‘From 8:15-8:30am I continued to eat breakfast’). A large emerging literature has shown that ongoing experiences are structured into events, whereby events are separated by event boundaries (Kurby & Zacks, 2008; Radvansky & Zacks, 2014; Zacks, Speer, Swallow, Braver, & Reynolds, 2007), and information within an event has greater overlap in shared context and content (Clewett & Davachi, 2017; Clewett, DuBrow, & Davachi, 2019; DuBrow & Davachi, 2014). In turn, this event structure influences memory organization (Clewett, Gasser, & Davachi, 2020; DuBrow & Davachi, 2013, 2014, 2016; Ezzyat & Davachi, 2011, 2014; Heusser, Ezzyat, Shiff, & Davachi, 2018; Pettijohn & Radvansky, 2018; Pettijohn, Thompson, Tamplin, Krawietz, & Radvansky, 2016; Radvansky, Krawietz, & Tamplin, 2011; Sols, DuBrow, Davachi, & Fuentemilla, 2017; Swallow, Zacks, & Abrams, 2009; Swallow et al., 2011; Zwaan, 1996), and subjective temporal estimates (Clewett et al., 2020; DuBrow & Davachi, 2013; Ezzyat & Davachi, 2014; Faber & Gennari, 2017; Lositsky et al., 2016).

In this way, event segmentation provides a window into the relationship between temporal representations and memory organization. Indeed, recently there has been increasing interest in the interaction between event segmentation, temporal information and memory organization (Clewett et al., 2019; Frank, Norman, Ranganath, Zacks, & Gershman, 2020; Radvansky & Zacks, 2017). However, most studies exploring the link between temporal information and event segmentation have queried temporal memory explicitly (e.g. DuBrow & Davachi, 2013, 2014; Ezzyat & Davachi, 2014; Faber & Gennari, 2017; Lositsky et al., 2016). By contrast, studies that have focused more narrowly on temporal information per se, have used more open-ended search tasks like free recall, in which participants recall as many items as possible from a just-presented list, in any order (e.g. Healey, Long, & Kahana, 2019; Sederberg, Miller, Howard, & Kahana, 2010). This leaves unclear how endogenous temporal information and event segmentation interact to organize memory during relatively unconstrained memory search tasks. Here we present a computational model which formalizes how event boundaries influence temporal information and memory representations. We verify novel predictions of this model using human behavior and neural activity, confirming the impact of event structure on temporal representations during memory encoding and retrieval.

Before turning to our novel approach and model predictions based on this three-way interaction of temporal representations, memory, and event segmentation, we begin by reviewing studies of the interaction between event segmentation and memory. On a behavioral level, there is a wealth of data to suggest that stimuli presented in the same event share stronger associations in long-term memory than stimuli presented in different events (DuBrow & Davachi, 2013, 2014, 2016; Ezzyat & Davachi, 2011, 2014; Heusser et al., 2018; Speer & Zacks, 2005; Zwaan, 1996). For instance, recognition memory for recently presented information is worse when an event boundary occurs between presentation and test (Swallow et al., 2009, 2011). Neural data corroborate these findings, as neural activity for pairs of stimuli from the same event exhibit stronger correlations than stimulus pairs from different events (Baldassano et al., 2017; DuBrow & Davachi, 2013, 2014; Ezzyat & Davachi, 2014; Hsieh, Gruber, Jenkins, & Ranganath, 2014; Lositsky et al., 2016; Schapiro, Roger, Cordova, Turk-Browne, & Botvinick, 2013). Further, brain activity in mnemonic brain regions (e.g. hippocampus) is greater during retrieval of items from another event rather than the current event (Swallow et al., 2011), suggesting that retrieval might be more effortful for information outside of the current event. Taken together, these results suggest that associations are weaker between memories separated by an event boundary, and further that overcoming such weakened associations may require more effortful retrieval.

Why might an event boundary weaken associations in memory? One account posits that event boundaries weaken temporal associations. In support of this account, stimuli separated by event boundaries are perceived as occurring farther apart in time than stimuli occurring in the same event (Ezzyat & Davachi, 2014; Faber & Gennari, 2017; Lositsky et al., 2016; Speer & Zacks, 2005; Zwaan, 1996). As an alternative account, stimuli separated by an event boundary may simply share fewer common features, independent of temporal associations. For instance, event boundaries may be caused by physical changes to the environment, such as a change in background scene (Zacks et al., 2007). Thus, weakened associations between information presented in different events may reflect the paucity of shared perceptual or categorical features, rather than weakened temporal associations. Few studies have distinguished between these accounts. Because most studies of event segmentation operationalize event boundaries as changes to the stimuli themselves, it is difficult to discern the unique contribution of temporal associations. Thus, this motivates our current study to examine the interaction between temporal associations, memory, and event segmentation. Before describing our approach, we first review other studies which provide some evidence that event boundaries weaken temporal associations, even in the absence of large changes in stimuli features.

In a series of studies with more controlled changes to stimuli between events, DuBrow and Davachi (2013, 2014) found that event boundaries influenced memory performance and memory representations. They defined an event as a sequence of presented stimuli from the same semantic category and with the same encoding task. In each list, they presented participants with sequences of items, switching back and forth between the two categories and tasks. Critically, they tested participants with pairs of items, where each pair contained items from the same category and task, but only a subset of pairs were from the same event. With these test stimuli, participants exhibited less accurate mnemonic judgments for item pairs across events than within event. This suggests that weakened associations across events are not completely a by-product of fewer shared stimulus features, which in turn suggests a role of temporal information. Further, DuBrow and Davachi (2014) found evidence suggesting that items within the same event, benefiting from strengthened associations, can support reinstatement of one another and their shared event information. In particular, participants were presented with two items previously studied with the same category and task, and determined which item was presented more recently. On a neural level, the category of the intervening items was decoded using whole brain multivariate pattern analysis (Norman, Polyn, Detre, & Haxby, 2006), and classifier performance predicted the category of these intervening items. On a behavioral level, DuBrow and Davachi (2014) posited that, if testing items from the same event evokes event-level reinstatement, then this reinstatement should facilitate memory recognition of other items from that event. Consistent with this hypothesis, participants recognized an item more quickly when it was preceded by a recency judgment of other same-event items. Taken together, these results reflect the strong associations between items within an event, and how such associations promote memory reinstatement of event-related information. Studies using television episodes, rather than discrete stimuli, have also found such neural evidence of event-level reinstatement (Baldassano et al., 2017; Chen et al., 2017; Zadbood, Chen, Leong, Norman, & Hasson, 2017). Nonetheless, in these studies, event boundaries were even more strongly defined by changes to stimulus features, and thus these results do not completely isolate the contribution of temporal similarity from stimulus similarity.

In a series of studies, Polyn, Norman, and Kahana (2009a, 2009b) examined the role of temporal information with even more consistent stimulus features across events. Although they did not frame their results in terms of event boundaries, like the DuBrow and Davachi studies, participants studied items with one of two encoding tasks, and thus a sequence of items with the same task can be operationalized as an event. Critically, to distinguish between event-level and temporal information, Polyn et al. (2009a) examined predictions of a computational model of episodic memory, the Context Maintenance and Retrieval (CMR) model. CMR assumes that two types of context are updated whenever an item is studied or retrieved: (a) temporal context, reflecting the surrounding temporal information of a given item; (b) event context, implemented experimentally as an encoding task. Polyn et al. (2009a) compared two variants of the CMR model: (1) one variant assumed that an event boundary evokes a change to event context only; (2) another variant assumed that an event boundary evokes a change to event context as well as a disruption to temporal context. The second CMR model variant made more accurate predictions of participants’ memory performance (Polyn et al., 2009a), suggesting that, even when accounting for differences between stimuli occurring in different events, an event boundary imposes a perceived shift or disruption in temporal information. These results underscore the critical role of temporal information to how events are structured and become represented in memory.

### The current study

Thus far, we have reviewed how event segmentation influences memory, and studies dissociating the contributions of temporal and nontemporal information to event boundaries and memory. Yet event segmentation has strong effects on temporal perception (Ezzyat & Davachi, 2014; Faber & Gennari, 2017; Lositsky et al., 2016; Speer & Zacks, 2005; Zwaan, 1996). Here we assert that an understanding of the interactions between event segmentation and memory remain incomplete without appreciating the role of temporal information. Specifically, it is critical to distinguish between the possibility that temporal representations are a defining feature of stimuli, and thus influenced by event boundaries, versus the possibility that temporal perception effects are a by-product of changes to other stimulus features. Distinguishing between these possibilities is not only important in the context of event segmentation, but more broadly may inform the role of temporal information to other perceptual and memory paradigms.

Critically, to our knowledge no research directly links the impact of event segmentation at study, including its impact on temporal disruption, to neural and behavioral measures of memory retrieval. Here we examined these relationships among memory behavior and a neural measure of temporal context (Folkerts, Rutishauser, & Howard, 2018; Howard, Viskontas, Shankar, & Fried, 2012; Manns, Howard, & Eichenbaum, 2007; Manning, Polyn, Baltuch, Litt, & Kahana, 2011). This neural measure allowed us to assess how temporal context states from study were reinstated during memory retrieval to influence behavior.

CMR provides an ideal testbed to examine the links between memory, temporal information, event cognition. CMR is a model of episodic memory sharing many assumptions with theories of event cognition (e.g. DuBrow & Davachi, 2014; Ezzyat & Davachi, 2014; Faber & Gennari, 2017; Frank et al., 2020; Lositsky et al., 2016; Swallow et al., 2009). We compared CMR predictions to data averaged across participants and we examined individual variability across participants. If temporal disruption underlies event segmentation and memory representations, then we expect (a) accurate predictions from the CMR model; (b) those participants exhibiting the largest disruption to temporal information to also exhibit the largest modulations during retrieval. To test these hypotheses, we present novel analyses of a neural correlate of temporal context, as posited by the CMR framework (Manning et al., 2011), as well as analyses of memory behavior which have been used to assess variants of the CMR model (Kahana, 1996; Lohnas & Kahana, 2014; Polyn et al., 2009a; Sederberg, Howard, & Kahana, 2008). We found that CMR predictions were upheld in averaged data, and participant variability was consistent across predicted measures. Our results clarify how event segmentation impacts temporal representations during memory encoding and retrieval, influencing perception and memory.

## Results

### Temporal context in control lists

We first assessed behavioral and neural measures of temporal context in control lists (Figure 1). These lists did not impose a strong event structure because participants performed the same (or no) encoding task for every studied item in each list (e.g., compare with Figure 3A). Thus, we used the control lists to test core assumptions regarding temporal context during study and retrieval. These form the foundation of our analyses assessing how temporal context is influenced by event segmentation.

**Figure 1:**
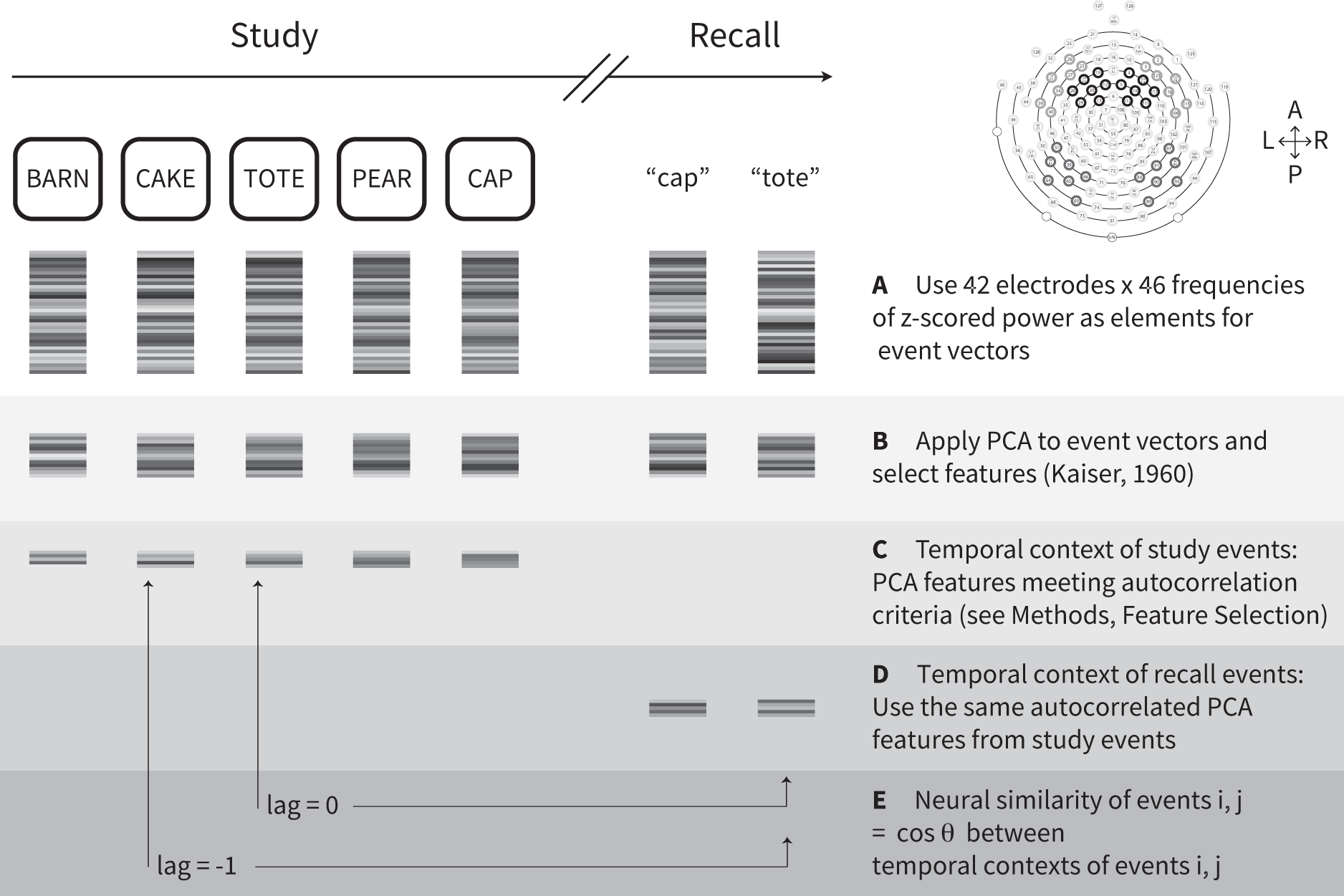
Calculating a neural measure of temporal context. Based on the core assumptions of retrieved context models, the temporal context state of a studied item should be a slowly changing representation of temporal context from earlier studied items; be reinstated if the item is recalled. **A.** We first calculated oscillatory power from electroencephalography (EEG) activity recorded for each studied item or recalled item in control lists. In the upper right panel, the 42 electrodes included in the event vectors are circled in dark gray on the electrode map. L = left, P = posterior, R = right, A = anterior. By applying principal components analysis (PCA), we selected features accounting for a significant amount of variance in the EEG recordings. **C.** To meet the first criterion of a slowly changing representation, we next determined which of the PCA features were autocorrelated across studied items. **D.** To verify the second criterion of a neural measure of temporal context, we next needed to examine this neural signature at recall. Thus, having established a slowly changing neural signature from study of selected PCA features, we then applied those same feature vectors from study events to the recall events. **E.** We assessed whether a studied item’s feature vectors were reinstated when the item was recalled, by calculating the neural similarity between each recalled item’s temporal context and temporal context states from study. Retrieved context models predict that the similarity between a recalled item’s retrieved temporal context and temporal contexts at study should be greater for items studied nearby in time, or smaller absolute lag, to the study position of the recalled item (see also Figure 2C,D).

### Evidence of temporal context in recall behavior

After studying each list, participants performed free recall, recalling as many items as possible from the just-studied list in any order. Despite the open-ended instructions, recall order tends to reflect the temporal order in which items were presented (Kahana, 1996; Kahana, Howard, & Polyn, 2008; Healey et al., 2019; Healey & Kahana, 2014; Ward, Tan, & Grenfell-Essam, 2010; Unsworth, Spillers, & Brewer, 2012). Contributions of temporal organization can be measured by calculating the probability of a recall transition between two items, based on their difference in serial positions at study and conditional on their availability (lag-CRP; Kahana, 1996). Figure 2B shows the lag-CRP from the control lists, demonstrating two ubiquitous and critical features of this function (Kahana et al., 2008). First, the lag-CRP tends to be greatest at smaller absolute lags, indicating the increased transition probability between items studied nearby each other. Second, the lag-CRP is asymmetric, with greater transition probability in the forward direction (positive lag) than the backward direction (negative lag).

**Figure 2:**
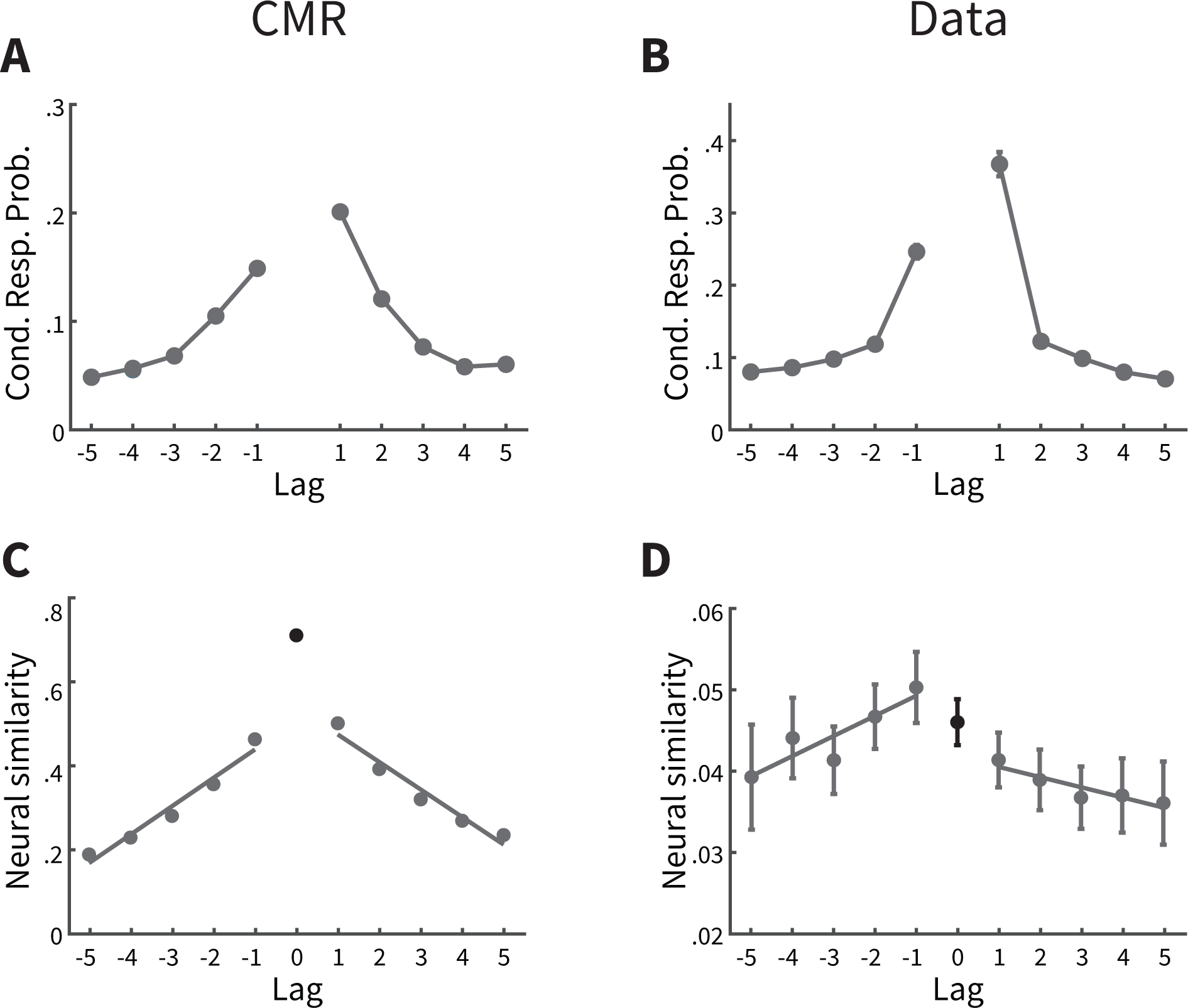
Behavioral and neural correlates of temporal context reinstatement in control lists. **A.** Predictions of the context maintenance and retrieval (CMR) model of recall transitions in control lists. The probability of making transitions between successive recalls is plotted as a function of lag, or difference in the serial positions of the successively recalled items. These response probabilities are determined conditional on which lags are available for recall. **B.** Conditional response probability as a function of lag in the behavioral data. **C**. Neural similarity between the temporal context state of a recalled item and the temporal contexts of its neighbors during study. Lag refers to the distance in serial position between two items from study (see Figure 1E). CMR predicts that temporal context states will be more similar between the recalled item and neighboring items from study. **D**. CMR’s predictions are upheld when measuring the neural measure of temporal context in participants’ data. Cond. Resp. Prob. = Conditional response probability. Error bars represent Loftus and Masson (1994) 95% confidence intervals.

Figure 2A presents CMR’s prediction of the lag-CRP and these two critical features. Rather than find a set of model parameters that best capture the phenomena of this data set, here we took a stricter approach by using a set of model parameters generated from another data set with a similar experimental design( (i.e., the best-fit parameters of the full model from Polyn et al. 2009, available at http://memory.psych.upenn.edu/CMR). From these simulations, CMR predicts the temporal contiguity effect, or tendency to recall items from smaller absolute lags. CMR predicts this effect due to its core assumptions that temporal context changes slowly with each studied item, and recall of an item leads to retrieval of its associated context states from study. Thus, when the current context cues recall of the next item, the just-recalled item’s context forms a part of this retrieval cue. As a result, CMR is more likely to recall items with shared temporal context states to the just-recalled item, including that item’s neighbors from study. CMR predicts the forward asymmetry in the lag-CRP because the context of a particular item *i* is incorporated into the context state of item *i* + 1, and thus a recalled item generally has a temporal context more similar to the items presented after it. Taken together, we interpret these behavioral recall dynamics as evidence for a role of temporal context in control lists.

### A neural signature of temporal context during study

According to CMR, the behavioral effects in Figure 2 rely on a temporal context representation which changes slowly with each studied item (Figure S1A). We assessed this prediction by defining an electrophysiological measure of temporal context consistent the definitions of Manning et al. (2011). Whereas Manning et al. (2011) examined temporal context with intracranial EEG, here we examined temporal context with scalp EEG, which unlike intracranial EEG is noninvasive. To our knowledge, our results provide the first evidence of a temporal context measure using scalp EEG.

The measure of temporal context was designed to meet several criteria consistent with CMR’s assumptions. First, we defined a vector of power values for each study event and recall event, where values were concatenated across a range of frequencies and included activity from electrodes implicated in mnemonic processing (Figure 1A; see Methods for full details Long, Burke, & Kahana, 2014; Long & Kahana, 2017; Weidemann, Mollison, & Kahana, 2009). We then applied principal components analysis to the matrix of power vectors across study and recall events, and excluded principal components that contributed a low level of variance in principal components space (Kaiser, 1960). Next, we quantified the extent to which each principal component changed slowly with each studied item, based on its autocorrelation (Equation 1; Figure 1C). If a principal component was not sufficiently autocorrelated across studied items, it was excluded because it did not meet the critical criterion of temporal context, to change slowly with each studied item. In this way, we calculated a set of autocorrelated feature vectors consistent with the notion of temporal context, separately for each participant and session. Our success in finding such autocorrelated feature vectors for 171/172 (99%) of participants attests to the validity of this approach (see also Figure S1B).

### Reinstatement of temporal context in control lists

We next verified CMR’s core prediction that temporal context is reinstated during recall. For each recalled item, we calculated the similarity between the temporal context state of that item when it was originally studied to the current temporal context state as the item was being recalled. In addition, we calculated the similarity between the current temporal context and the temporal contexts of the item’s neighbors from study (Figure 1E). According to CMR, because an item’s recall leads to reinstatement of temporal context state from study, then the neural similarity between the current context and an item’s context from study should reflect the temporal history of studied items, such that items with smaller absolute lags should have greater similarity (Figure 2C). We assessed CMR’s prediction with the autocorrelated feature vectors, our alleged neural measure of temporal context. Specifically, for each recalled item, we calculated the neural similarity between the recalled item’s feature vector to both the feature vectors from study of itself (lag = 0) and to its neighbors of lag *∈ {−*5*, −*4*, …,* 4, 5*}*, for those items not yet recalled (Figure 2D; also see Materials and Methods). For negative lags (i.e., the similarity between an item and the items studied before it), CMR predicts that neural similarity should increase with study-recall lag. This is because a recalled item’s retrieved temporal context should have greater overlap in temporal context, i.e. greater neural similarity, with other items studied nearby in time to that item. This prediction is critical to distinguish the retrieval of *context* information, as predicted in CMR, from the retrieval of *content*, or item, information (Manning et al., 2011). To test this prediction, we compared the neural similarity between at lag = -1 to the neural similarity at more distant lags -3 to -5. Consistent with CMR’s prediction, neural similarity was significantly greater at lag =-1 than the average neural similarity at lags -3 to -5 (*M* = 0.009*, SD* = 0.039*, t*(169) = 2.95*, CI* = [0.0029, 0.0146]*, p* = 0.004).

We also evaluated neural similarity at positive lags, predicting that neural similarity should decrease with lag in the forward direction. Following the logic with negative lags, CMR predicts that context states of items studied nearby in time should share more temporal context, thus leading to greater neural similarity. Paralleling the test of neural similarity with negative lag, we compared the neural similarity at lag = +1 to the neural similarity at lags 3 to 5, and found that neural similarity was greater at lag = +1 (*M* = 0.005*, SD* = 0.031*, t*(169) = 1.99*, CI* = [0.0000, 0.0095]*, p* = 0.048).

This result is not only consistent with CMR’s prediction, but also helps to rule out the possibility that our neural measure of temporal context only reflects autocorrelated noise (Manning et al., 2011).

### The influence of event boundaries on temporal context representations

Having established neural and behavioral measures of temporal context in control lists, we next turned to the critical analyses of the influence of event boundaries on temporal context states. Specifically, CMR predicts that the similarity in temporal context between neighboring studied items should be less if those items are separated by an event boundary, due to a disruption in temporal context. Further, CMR predicts that the state of temporal context, incorporating the disruption to temporal context during study, should be reinstated during recall. We examined these predictions in two-task lists, where participants performed 1 of 2 semantic encoding tasks with each studied item, switching back and forth between the 2 tasks throughout the list (Figure 3A). In this way, an event is operationalized as a sequence of items presented with the same task, where task was indicated to the participant by the color, font and case of the word. We define an event boundary defined as a change in encoding tasks. Thus, we term a *boundary* item as the first item presented with the changed encoding task, and a *preboundary* item as the final item in an event before the task switch.

**Figure 3:**
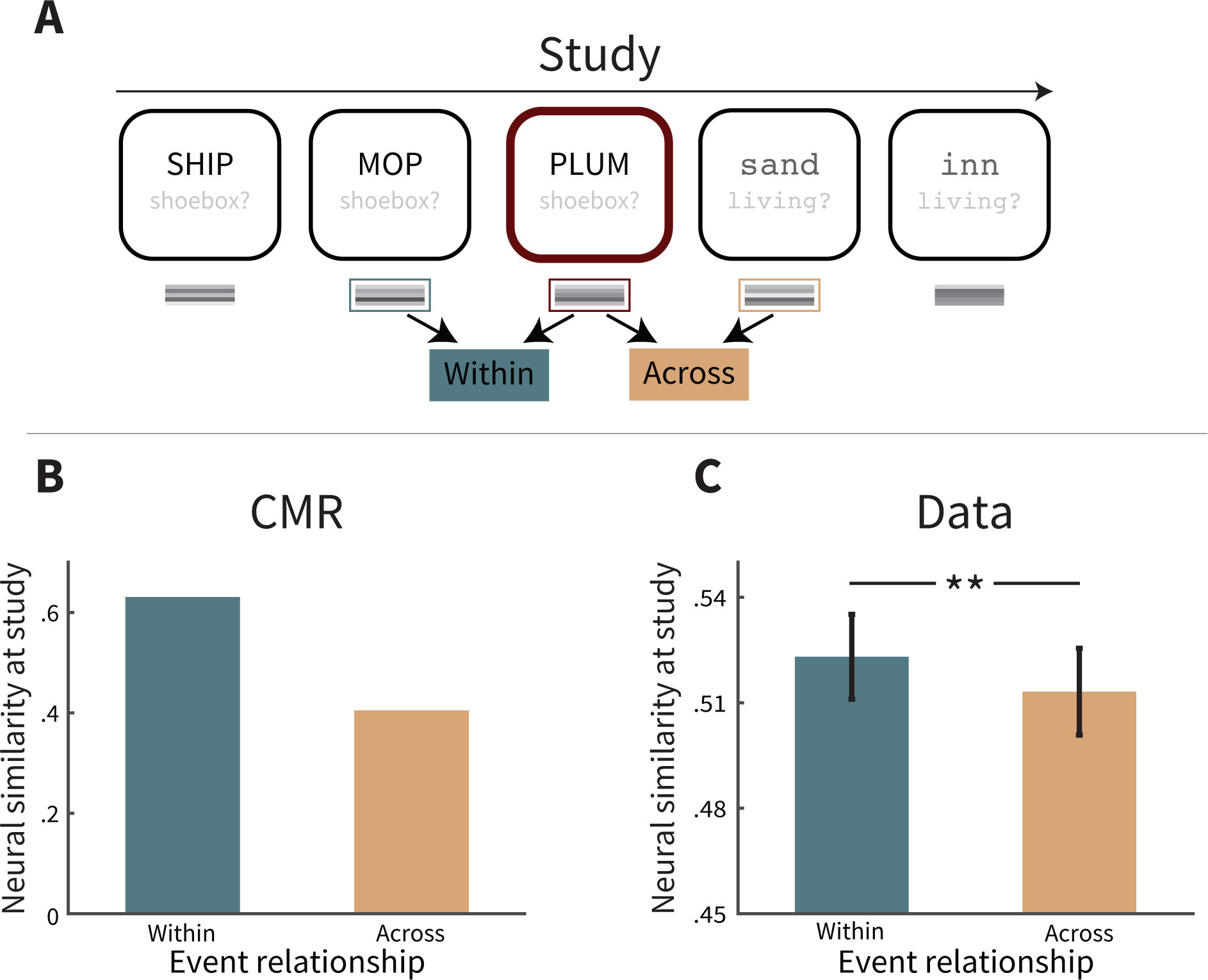
Modulation of temporal context by event boundaries during study. **A.** In two-task lists, participants perform one of two encoding tasks with each presented word (Shoebox task or Living task); a sequence of words with the same task is assumed to comprise an event, and the change in task is assumed to form an event boundary. Here the sample items are shown to calculate neural similarity for the neighbors of a preboundary item, as one of its neighbors (the preceding item) was presented within the same event (Within), and its other neighbor (the following item) was presented across a different event (Across). Task text is for illustrative purposes only; to participants this was implicit from the color, font and case of the word. **B.** Collapsed across preboundary and boundary items, CMR predicts that neural similarity is greater between two neighboring items within the same event than two items across different events. **C**. Collapsed across preboundary and boundary items, neural similarity is greater between two neighboring items within the same event than two items across different events. Error bars represent *±*1 standard error of the mean. ** *p < .*01.

In the experimental data, we calculated temporal context for each item in each two-task list using the feature vectors from the control lists (Figure 1). We excluded the two-task lists when calculating temporal context, so that any context modulations observed in these lists could not be explained by variance in the lists themselves. This approach shares similarities to cross-validation, whereby a subset of the data is used for training and its complement is used to test the data. In particular, the results reported below cannot be explained based on contributions of within-list task switches to temporal context, as such switches did not exist in the lists we used to define temporal context. Nonetheless, we first verified that the temporal context features, defined in control lists, still maintained the critical property of autocorrelation in the two-task lists. We calculated neural similarity during study between items from the same event, and found that neural similarity was significantly greater for items with lag = 1 than lags 3 to 5 (*M* = 0.070*, SD* = 0.064*, t*(169) = 14.41*, CI* = [0.0606, 0.0799]*, p <* 0.0001; Figure S1B). With respect to model simulations, temporal context states in CMR are just as easily measured in the control or two-task lists based on model equations (Figure S1A; also see Appendix). Having defined event structure and temporal context in the two-task lists, we now assess CMR predictions in the experimental data.

### Event boundaries modulate temporal context during study

CMR assumes that an event boundary leads to a disruption in temporal context, making temporal context after the event boundary less similar to the prior temporal context state. Thus, holding lag constant, the neural similarity in temporal context between two items should be less when those items are separated by an event boundary. CMR predicts that neural similarity should be less across boundaries at any lag, yet because context similarity also decreases with lag, at larger lags this difference becomes more subtle. Thus, CMR predicts the most salient influence of event boundaries for neighboring pairs of items, and here we examine this stricter test of CMR’s predictions at study lag = 1. Figure 3B shows CMR’s prediction of the neural similarity between pairs of successive items that border an event boundary, as a function of being presented in the same event or different events.

We tested CMR’s prediction by calculating the neural similarity between temporal context feature vectors of successive items bordering an event boundary during study (Figure 3C). We found that neural similarity was greater between item pairs studied within the same event than item pairs studied across different events (*M* = 0.009*, SD* = 0.042*, t*(169) = 2.81*, CI* = [0.0027, 0.0154]*, p* = 0.006). This result is consistent with CMR’s underlying assumption that there is a disruption to temporal context at the event boundary, thus leading to reduced temporal similarity of items separated by an event boundary. This also suggests that, for items separated by an event boundary, their weakened neural similarity may reflect their weakened temporal associations.

### Reinstatement of event modulations to temporal context

We next queried temporal context representations during recall, motivated by CMR’s assumption that recall of an item evokes retrieval of its context states from study. In the two-task lists, the retrieved context includes the temporal context modulated by event boundaries. Thus, CMR predicts that if a preboundary or boundary item is recalled, then the retrieved temporal context of the recalled item should show greater similarity with its within-event neighbor from study in comparison its across-event neighbor from study (Figure 4A,B). However, the predicted difference in neural similarity is more subtle during recall than during study (i.e., compare to Figure 3B). In retrieved context models such as CMR, the extent of context reinstatement for each item is defined by a parameter ranging from 0 to 1, with 0 indicating no context reinstatement and 1 indicating perfect reinstatement. If context reinstatement was perfect, then the predictions of study and recall would be the same. To best account for human behavior, the context reinstatement parameter is less than perfect (here, set to .510; see Table 1). As a result, the difference for within-event similarity vs. across-event similarity during recall is less than (a perfect reflection of) the difference in context from study.

**Figure 4:**
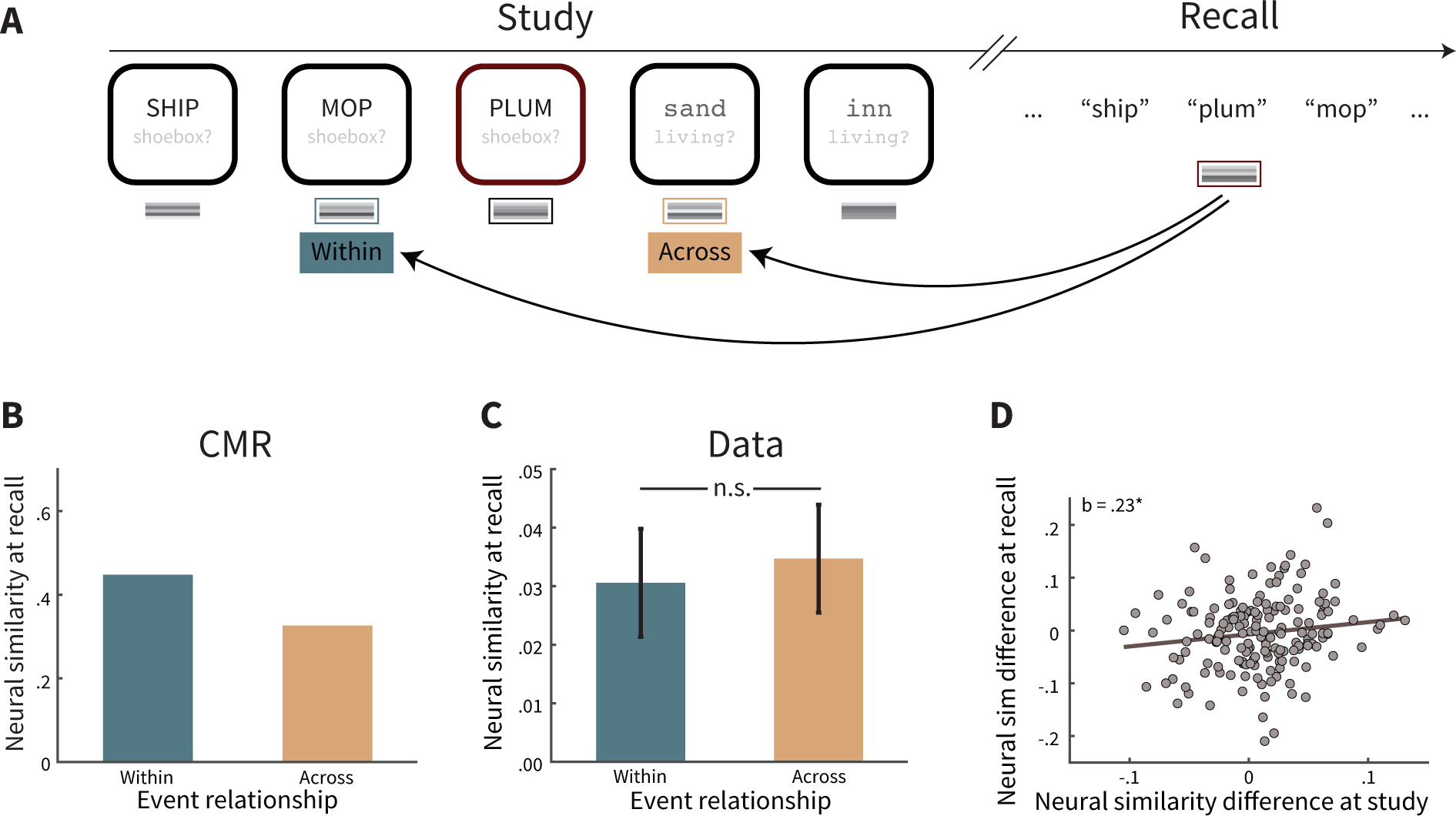
Neural similarity during recall. **A.** Calculation of the neural similarity of the current state of context after recall of a preboundary item with its associated neighbors at study: both the preceding neighbor presented with the same task, and thus within the same event (Within), and with its subsequent neighbor across a different event (Across). Task text is for illustrative purposes only; to participants this was implicit from the color, font and case of the word. **B.** CMR predicts that the retrieval of an item bordering an event boundary (e.g. ‘plum’ in A) leads to retrieval of that item’s temporal context states from study, including the disruption to temporal context caused by the event boundary. Thus, the current state of temporal context—which incorporates the item’s retrieved temporal context—should be more similar to the context of adjacent studied item within the same event (e.g. ‘mop’) than the adjacent studied item from a different event (e.g. ‘sand’). However, the difference by event relationship is more subtle than during study (compare with 3B). **C**. Mean neural similarity in the behavioral data was not significant by event relationship. Error bars represent ±1 standard error of the mean. **D.** Participants exhibiting greater disruption to temporal context during study also exhibit a greater reinstatement of disruption in temporal context during recall. * *p < .*05 (one-tailed).

**Table 1:**
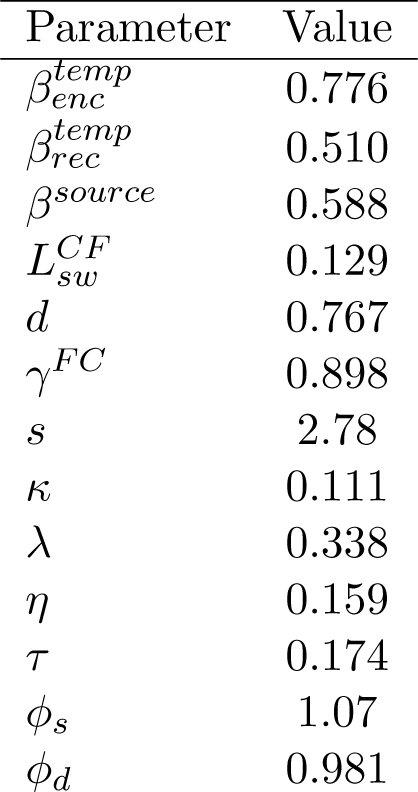
Best-fit parameters from the full model of Polyn et al. 2009, determined using a genetic algorithm fitting technique

| Parameter | Value |
| --- | --- |
| $\beta_{enc}^{temp}$ | 0.776 |
| $\beta_{rec}^{temp}$ | 0.510 |
| $\beta^{source}$ | 0.588 |
| $L_{sw}^{CF}$ | 0.129 |
| $d$ | 0.767 |
| $\gamma^{FC}$ | 0.898 |
| $s$ | 2.78 |
| $\kappa$ | 0.111 |
| $\lambda$ | 0.338 |
| $\eta$ | 0.159 |
| $\tau$ | 0.174 |
| $\phi_s$ | 1.07 |
| $\phi_d$ | 0.981 |

We next assessed CMR’s prediction in participant EEG data for recall of boundary items and preboundary items (Figure 4C). Neural similarity was not significantly different for a recalled item if the similarity was calculated with its studied neighbor from the same versus a different event (*M* = *−*0.004*, SD* = 0.070*, t*(169) = *−*0.78*, CI* = [*−*0.0147, 0.0064]*, p* = 0.439). Yet this is not entirely inconsistent with CMR’s prediction that the difference in similarity reinstated during recall is less than the neural similarity difference at study. To further probe whether this nonsignificant difference in neural similarity might reflect a meaningful signal in temporal context, we examined the across-participant variability in temporal context reinstatement. In particular, we hypothesized that those participants exhibiting greater modulations in temporal context during study should also exhibit greater reinstatement of those modulations during recall, even if the mean difference in temporal context was not significant. To test this hypothesis, we calculated each participant’s difference in neural similarity at study (within vs. across event in Figure 3C), and correlated this with each participant’s neural similarity difference at recall (Figure 4C). Using robust regression, we found that these two neural similarity measures were correlated (*N* = 170*, b* = 0.23, one-tailed *p* = 0.032; Figure 4D). This suggests that, if a participant experiences the task changes as more salient disruptions to temporal information associated with items, then that participant also reinstates such temporal disruptions when recalling those items. Thus, this result is consistent with our hypothesis that the disruptions to temporal context by event boundaries from study were reinstated during recall.

### Event modulations to temporal context from study influence recall behavior

We next examined a novel prediction of CMR concerning the impact of event boundaries on free recall behavior, in particular recall transitions. This prediction builds on previous findings establishing that much of the variability of recall transitions in free recall can be explained by the temporal relationships between studied items, as participants are more likely to transition between items presented nearby on the study list (Figure 2B; Healey et al., 2019; Kahana, 1996, 2012). CMR assumes that temporal context drives this temporal organization (Figure 2A), and so CMR also assumes that recall transitions should be modulated by temporal disruptions imposed by event boundaries. As shown in Figure 5A, CMR predicts that recall transitions *from a preboundary item* should be less likely to the item at lag = +1 in the two-task lists, as compared to transitions of lag = +1 in the control lists (Figure 2A). This striking prediction stands in contrast to the forward asymmetry usually seen in free recall (Healey et al., 2019; Kahana, 1996, 2012). Yet, according to CMR, an event boundary disrupts temporal context between the preboundary item and the next item (at lag = +1), and so these items do not overlap as much in their temporal context states. As a result, when the preboundary item is recalled and its temporal context is reinstated, the retrieval cue incorporating this context will overlap less with the context of the lag = +1 item. Thus, this state of context does not promote recall of the lag = +1 item as strongly as in a control list. Further, more temporal context is shared between the lag = -1 item and the preboundary item, because these items are from the same event. Thus, CMR predicts that transitions from a preboundary item to the item studied before it, the lag = -1 item, is more likely when compared to control lists or even to the lag = +1 item.

**Figure 5:**
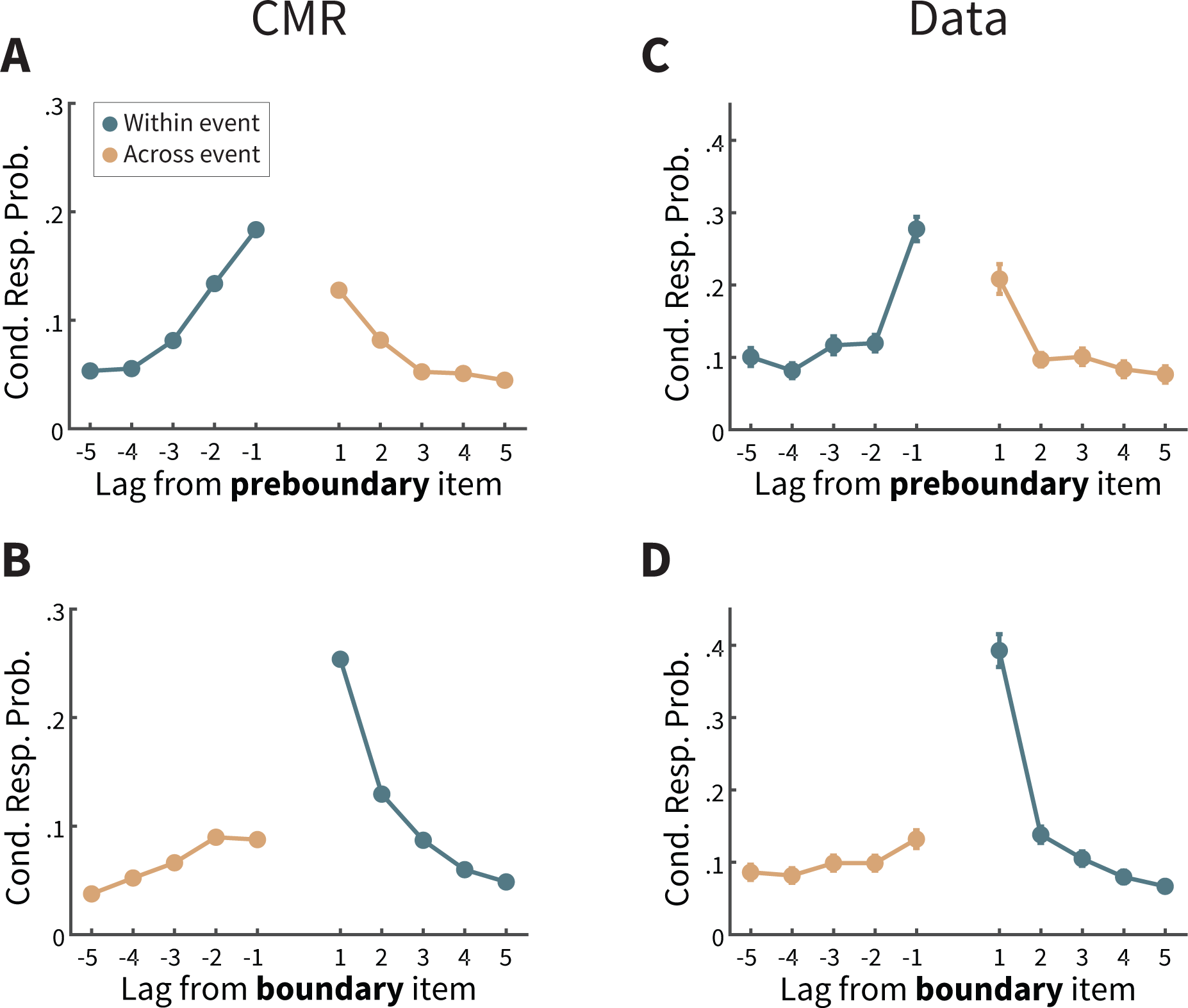
Recall transitions in two-task lists. **A.** The context maintenance and retrieval (CMR) model predicts that transitions from a preboundary item are more likely to be other items within the same event (teal lines) than to items in the following event (orange lines), in contrast to the established bias to make forward transitions (see Figure 2B). **B.** CMR predicts that transitions from a boundary item are more likely to items in the same event (teal lines) than to items in the preceding event (orange lines), leading to an exaggerated tendency to make forward transitions. **C,D.** Consistent with CMR predictions, participants are more likely to recall items presented in the same event (teal lines) than the neighboring event (orange lines). Cond. Resp. Prob. = Conditional response probability. Error bars represent Loftus and Masson (1994) 95% confidence intervals.

In a complementary way, CMR predicts that recall transitions *from boundary items* are modulated as well (Figure 5B). A transition from a boundary item to its neighbor at lag = +1 should be more likely than in control lists, because these items share both event information and temporal information. Following similar logic, CMR also predicts that a transition from a boundary item should be less likely to the item at lag =-1, because such items were presented in a different event and thus share less temporal context with the just-recalled boundary item.

Next, we examined whether CMR’s predictions were upheld in participants’ data. To assess these effects statistically, we defined the *temporal modulation score* as the difference in lag-CRP values at *|* lag *|* =1 for transitions made within event minus transitions made across event. (Thus, for preboundary items this score is defined as CRP values at lag = -1 minus those at lag = +1; for boundary item this score is defined as CRP values at lag = +1 minus those at lag = -1). As a baseline, we compared this value to the lag-CRP values at the same lags from the control lists. We found that the distribution of temporal modulation scores from preboundary items was significantly greater in two-task lists than matched lags in control lists (*M* = 0.190*, SD* = 0.154*, t*(169) = 16.09*, CI* = [0.1670, 0.2137]*, p < .*0001). Qualitatively, the lag-CRP in the experimental data (Figure 5C) exhibits a similar pattern to CMR’s prediction, with larger values for transition probabilities for negative lags over positive lags. In addition, the temporal modulation scores from boundary items were also significantly greater in two-task lists than matched lags in control lists (Figure 5D; (*M* = 0.139*, SD* = 0.166*, t*(169) = 10.97*, CI* = [0.1144, 0.1646]*, p < .*0001). Thus, the recall transitions in two-task lists of the participants’ data are consistent with CMR’s assumption that event boundaries disrupt temporal representations, to promote transitions between items within the same event.

### Relating neural temporal context to recall behavior

Having established measures suggestive of the modulation of event boundaries on temporal context — neurally and behaviorally — we next asked whether the neural measures predicted the behavioral measures. According to CMR, both the neural modulation and the behavioral modulation should be greater when there is a greater disruption to temporal context by event information. Although CMR predicts average data, the extent of disruption may vary by participant. If this were the case, then those participants exhibiting a greater modulation by event boundary in their neural similarity difference should exhibit a greater modulation by event boundaries in their recall transitions. In other words, if each participant’s neural measures (Figure 4) and behavioral measures (Figure 5) both reflected modulations of temporal context by event boundaries, and if such information influenced memory, then the neural and behavioral measures should be correlated across participants.

We defined a participant’s *neural* modulation by temporal context as the difference in neural similarity for within-event transitions versus across event transitions (i.e., the difference in the two bars plotted in Figure 4). We defined a participant’s *behavioral* modulation by temporal context by summing the temporal modulation scores of preboundary and boundary items. Here we leveraged the variability across participants in the extent to which event boundaries modulate their temporal context states, and we predicted a positive correlation between neural modulation and temporal modulation. Again using robust regression to account for outliers, we found that across participants these two measures were correlated (Figure 6, *N* = 170*, b* = .41, one-tailed *p* = .045). This significant correlation indicates that variance in the temporal modulation scores can be explained by the neural temporal context measure. Although there is variability across participants in the extent to which event boundaries modulate their behavioral and neural activity, the significant correlation across participants suggests that a disruption to temporal context may underlie both effects. This correlation also argues against the possibility that the significant differences in the temporal modulation scores simply reflect recall organization based on shared event information and shared encoding task, rather than shared temporal context. If temporal context did not contribute to recall transitions, then we would not expect recall behavior to correlate with the neural measure of temporal context. Taken together, these results implicate the importance of temporal context for memory organization, and the influence of event structure on temporal context during encoding and retrieval.

**Figure 6:**
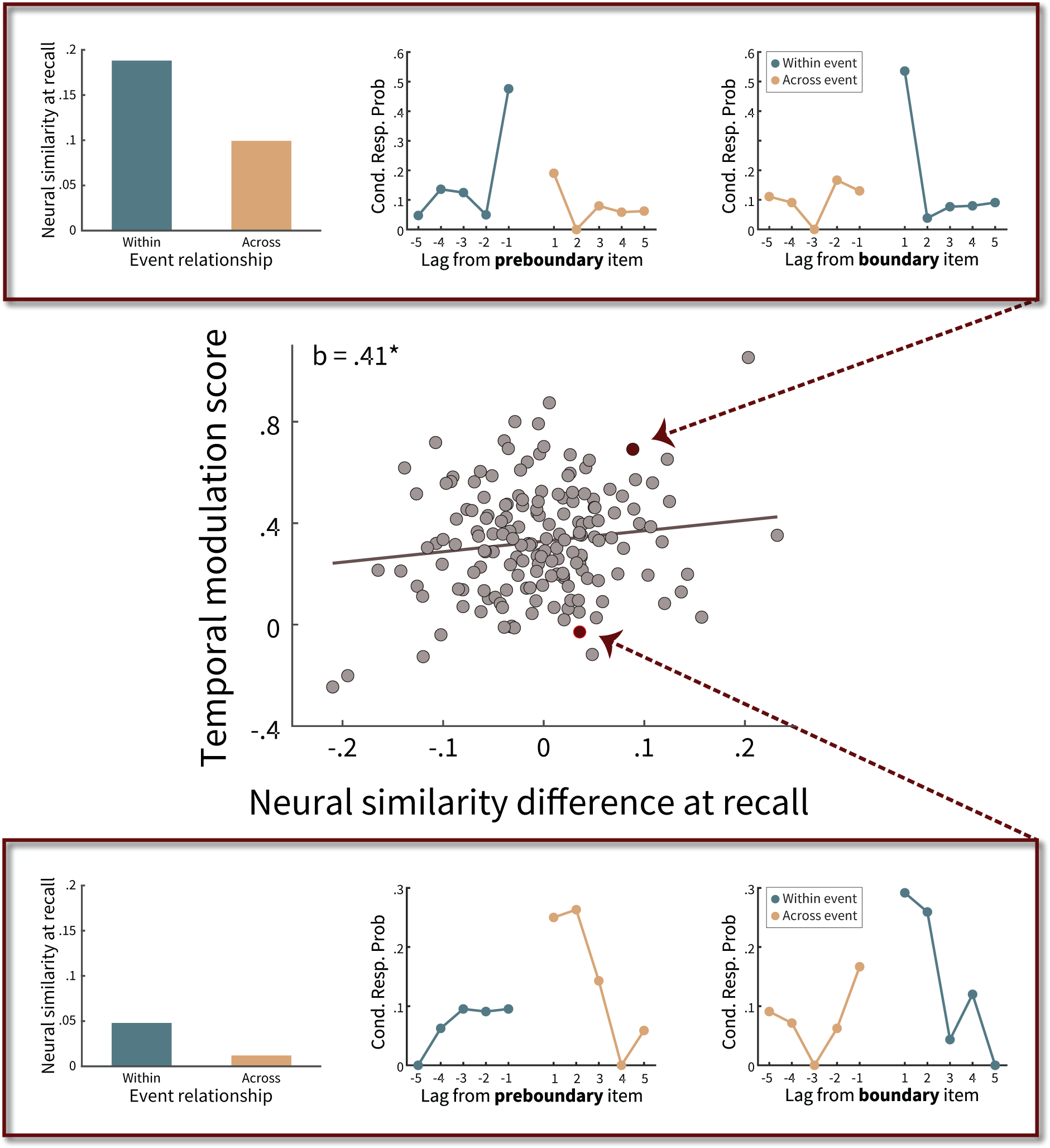
Influence of event boundaries on neural and behavioral measures of temporal context. In this correlation plot, each dot corresponds to a participant. The x-axis reflects the neural measure of event boundary modulation on temporal context; the y-axis reflects a behavioral measure of event boundary modulation on temporal context (see text for details). The top and bottom panels show for two participants the neural similarity at recall, used for calculating the x-axis, and conditional response probability (Cond. Resp. Prob.) as a function of lag, used for calculating the y-axis. Top panel: This participant has a high neural similarity difference at recall and a high temporal modulation score. Bottom panel: This participant has a low neural similarity difference at recall and a low temporal modulation score, as recall transitions are similar irrespective of whether the transition is from a preboundary item (bottom middle panel) or a boundary item (bottom right panel).

## Discussion

Human experiences and memories are not perfect reflections of the external environment. How differences emerge between the objective environment and internal experience remains a broad yet fundamental question in cognitive neuroscience. Such differences can impact perception of information in the moment, as well as how the information becomes represented in memory. Appreciating the interactions between ongoing perception and subsequent memory provides insight into both processes (Clewett et al., 2019; Zacks et al., 2007). Here we sought to link the influence of event segmentation on temporal representations and memory performance, by recording EEG as participants took part in free recall sessions. To discern the unique contribution of temporal information to event segmentation and memory organization, we tested neural and behavioral predictions of a computational cognitive model.

Our measure of temporal information differs from most other studies relating event segmentation and temporal representations, which ask participants explicitly to make temporal judgments (Ezzyat & Davachi, 2014; Faber & Gennari, 2017; Lositsky et al., 2016). Instead, we measured a neural correlate of temporal context with EEG. This enabled us to assess temporal perception both prospectively and retrospectively, and allowed us to query how temporal information influences memory dynamics even when such information is not as critical to task performance. The calculation of this temporal measure was motivated by retrieved context models such as CMR. These models assume that temporal context changes slowly with each studied item, and that an item’s temporal context state is retrieved when the item is recalled (Figure 1; Howard & Kahana, 2002; Manning et al., 2011). We first established this measure of temporal context in control lists, demonstrating for the first time a neural correlate of temporal context in scalp EEG (Figure 2).

Here we operationalized event boundaries with minimal changes to presented information: a change in the encoding task performed with each presented item, where the task was indicated by the color, font and case of the item. Event boundaries are often operationalized by more salient changes, such as a change in the semantic category of presented information (e.g., DuBrow & Davachi, 2013, 2014; Ezzyat & Davachi, 2014), features of the presented information such as size, location or color (e.g., Faber & Gennari, 2015, 2017; Heusser, Poppel, Ezzyat, & Davachi, 2016; Heusser et al., 2018; Lositsky et al., 2016; Radvansky & Copeland, 2006), as well as the predictability of upcoming information (e.g., Zacks et al., 2001). By contrast, we minimized stimulus changes between events to better isolate the contribution of temporal information to event structure.

With our conservative definition of event boundaries, we first verified that EEG activity can provide a neural measure of temporal context. We next verified several CMR predictions regarding the influence of event boundaries on temporal information. First, we found that neighboring items have reduced neural similarity in temporal context when separated by an event boundary (Figure 3). These results, measuring EEG activity extracted from electrodes across mnemonic regions (Long et al., 2014; Long & Kahana, 2017; Weidemann et al., 2009), are consistent with prior work finding reduced neural similarity across events in medial temporal lobe regions, including the hippocampus (Ezzyat & Davachi, 2014) and entorhinal cortex (Lositsky et al., 2016). Critically, we examined how temporal modulations during study influenced neural and behavioral activity during memory retrieval. While freely recalling list items, participants exhibited neural activity consistent with reinstatement of temporal context from study. In control lists, the neural temporal context of a recalled item was most similar to the temporal context of its neighbors from study (Figure 2D), consistent with predictions of the retrieved context model framework (Figure 2C). In addition, in the lists with two tasks and event structure, we found evidence that participants reinstated temporal context states from study, including disruption of temporal context disruptions between events. Specifically, participants exhibiting a larger decrease in temporal context similarity across events during study also exhibited a greater decrease in this measure during recall (Figure 4D). This provides support for temporal context reinstatement during recall, as those participants more influenced by the disruption of temporal information during study also reinstates such information during memory retrieval. Reinstatement of event structure, as reflected in temporal context, is consistent with prior evidence of event-related reinstatement during memory retrieval (Baldassano et al., 2017; Chen et al., 2017; DuBrow & Davachi, 2014; Zadbood et al., 2017). Yet it is noteworthy that the free recall test does not explicitly ask participants to remember this temporal information or the event structure. Thus, our results suggest that recall of an item automatically evokes retrieval temporal context states from study, consistent with CMR’s assumption.

Patterns of recall behavior, and their relation to neural activity, also attested to the influence of these neural temporal context states on memory organization. Although in a free recall task, participants may recall items in any order, recall order in two-task lists reflected the influence of event segmentation. Specifically, recall transitions were less likely between items studied in different events than items studied in the same event (Figure 5C,D), predictions that fall out naturally from CMR (Figure 5A,B) because event boundaries weaken memory associations by disrupting temporal representations. This pattern of behavior was most striking for successive recalls between neighboring items (i.e. lag = *±*1), and is consistent with findings of weakened memory associations between items across events versus within events (Baldassano et al., 2017; DuBrow & Davachi, 2013, 2014; Swallow et al., 2009, 2011). From the viewpoint of CMR, a participant exhibiting a greater difference in neural similarity during study experienced larger disruptions to temporal context, which should manifest during the recall test in both neural activity and memory performance. We found evidence of these relationships in mean participant data, as well as in across-participant variability. Future work remains to characterize how such changes vary by participant and by individual event, as well as whether more subtle changes to context might be better inferred as a shift in, rather than a disruption to, context (DuBrow, Rouhani, Niv, & Norman, 2017). Nonetheless, to our knowledge, this is one of the first studies to link directly, through neural activity and behavior, how event structure can influence temporal representations to impact memory performance.

Our neural measure of context, including EEG activity in mnemonic regions in the frontal and temporal lobes (Long et al., 2014; Long & Kahana, 2017; Weidemann et al., 2009), relates to prior studies of neural activity in episodic memory and event segmentation. More broadly, the medial temporal lobe has been implicated in episodic memory (Aggleton & Brown, 1999; Davachi, 2006; Eichenbaum, 2004; Fernandez et al., 1999; Goyal et al., 2018; Sugar & Moser, 2019) and temporal representations (Eichenbaum, 2014; MacDonald, Lepage, Eden, & Eichenbaum, 2011; Tsao et al., 2018). At the same time, the hippocampus exhibits reinstatement of event-level information in particular (Baldassano et al., 2017; DuBrow & Davachi, 2014), possibly supported by greater hippocampal activity after an event boundary (Baldassano et al., 2017; Sols et al., 2017). Further, the hippocampus is critical for the binding of episodic features, including those within an event (Davachi, 2004; Pacheco Estefan et al., 2019; Heusser et al., 2016; Staresina & Davachi, 2006, 2009; Richmond & Zacks, 2017). Regions of the frontal lobe are also critical for episodic memory (Blumenfeld, Parks, Yonelinas, & Ranganath, 2011; Jenkins & Ranganath, 2010; Hanslmayr & Staudigl, 2014; Long et al., 2014; McAndrews & Milner, 1991; Paller & Wagner, 2002) including free recall (Long, Oztekin, & Badre, 2010; Sederberg et al., 2007; Staresina & Davachi, 2006), as well as cognitive control processes exhibited in event segmentation (Baldassano et al., 2017; Chen et al., 2017; DuBrow & Davachi, 2016; Ezzyat & Davachi, 2014; Kurby & Zacks, 2008; Sols et al., 2017; Zacks et al., 2001). Thus, our results build on prior work implicating episodic memory regions in event segmentation, reflecting how they support encoding and retrieval of event information and influence memory organization.

Although we considered predictions of the CMR model, CMR’s predictions are consistent with other current cognitive model frameworks which segment events and make predictions of memory. However, unlike other models of event segmentation, CMR cannot infer event structure but needs to be provided the event structure explicitly (e.g. Radvansky, 2012; Zacks et al., 2007). Despite the different objectives of these types of models, CMR assumes event boundaries cause a disruption to memory associations, a similar assumption as established accounts of event processing, such as Event Segmentation Theory and the Event Horizon Model (Kurby & Zacks, 2008; Radvansky, 2012; Radvansky & Zacks, 2017; Zacks et al., 2001). Thus, our findings may be explained by the Event Horizon Model framework, which embodies Event Segmentation Theory. According to this framework, a current event model is held in working memory, and each event boundary updates the model. As a result, it is more difficult to retrieve information outside of the current event or the currently retrieved event (Radvansky & Zacks, 2017; Swallow et al., 2011). Thus, the Event Horizon Model should predict the decrease in recall transitions between neighboring items from different events. However, development of the Event Horizon Model has focused primarily on information repeated across events, whereas in the current study each event was comprised of unique novel items. Currently this model does not make quantitative predictions, so it remains to be fully developed to accommodate a stronger role of temporal information. Recently, Frank et al. (2020) presented Structured Event Memory (SEM), a computational model of event cognition. Like CMR, SEM can incorporate event structure during study to predict memory performance. Unlike CMR, SEM can infer event structure across a range of naturalistic stimuli, and can predict memory performance based on the inferred structure. Although CMR needs to be provided the event structure explicitly, in a list-learning paradigm as in the current study, SEM assumes that event structure is inferred correctly based on encoding tasks. Yet even if both models were provided with the event structure of the current study, SEM does not include explicit representations of temporal information nor has it been applied to free recall data. However, it would be an intriguing direction to examine how SEM might account for the effects presented here, in the absence of temporal information. In a complementary way, CMR may be a suitable framework to extend by incorporating more complex stimulus features and event structures. Such an extension would build upon CMR simulations accounting for the neural correlates of task representations (Morton et al., 2013) and temporal representations (Kragel, Morton, & Polyn, 2015; Manning et al., 2011). We have begun this process by presenting CMR predictions which integrate these two types of representations on a neural and behavioral level. Such predictions capture how objective temporal and event information interact to influence memory representations and retrieval.

An underlying objective of cognitive neuroscience and psychology concerns the transformation of external, objective environment into internal, subjective experience. This transformation begins during initial perception, and then influences how information is represented in, and retrieved from, memory. Although the relationships between temporal perception and episodic memory can complicate characterizations of their interactions, neural activity and computational models provide informative insight beyond behavior alone. Leveraging these methodological tools showed that event segmentation impacts temporal representations during initial perception and memory encoding. In turn, temporal representations influenced memory retrieval processing. Taken together, these results reveal how temporal information structures perception and memory, underscoring its importance in episodic memory and human cognition.

## Materials and Methods

### Participants

The data reported here are from the Penn Electrophysiology of Encoding and Retrieval Study (PEERS), which involved three subsequently administered multi-session experiments. PEERS aims to assemble a large database on the electrophysiological correlates of memory encoding and retrieval. The present study considered the 172 younger adults (age 17–30) who completed Experiment 1 of PEERS. Behavioral data is available at http://memory.psych.upenn.edu/files/PEERS.data.tgz and electrophysiology data is available at http://memory.psych.upenn.edu/mediawiki/ index.php?title=Data Request&paper=WeidKaha16.

Participants were recruited through a two–stage process. First, we recruited righthanded native English speakers for a single session to introduce participants to EEG recordings and the free recall task. Participants who did not make an excess of eye movements during item presentation epochs of the introductory session and had a recall probability of less than 0.8 were invited to participate in the full study. Approximately half of the participants recruited for the preliminary session qualified for, and agreed to participate in, the full study. Participants were consented according the University of Pennsylvania’s IRB protocol and were compensated for their participation.

One participant was excluded for not having a neural measure of temporal context in any session (see definitions below and Figure 1), and another was excluded from all behavioral and neural analyses for making too few (*<* 10) critical recalls (see Figure 4 and surrounding text). This participant had 7 such observations in total, whereas the next fewest participants had 15. For this latter participant, because most analyses include recall behavior, for consistency we excluded this participant from all, rather than just the recall, analyses.

### Experiment design

Participants completed 6 sessions each with 16 free recall lists. For each list, 16 words were presented one at a time on a computer screen followed by an immediate free recall test. Based on our criteria of only including sessions with autocorrelated feature vectors (see Feature selection), 5 participants had 2 included sessions, 8 participants had 3 included sessions, 24 participants had 4 included sessions, 50 had 5 included sessions, and 83 had 6 included sessions. Additional memory tests were administered in each session after immediate free recall of the final list. However, we do not report results from those tests so omit further detail about them.

Each word was accompanied by a cue to perform one of two judgment tasks (“Will this item fit into a shoebox?” or “Does this word refer to something living or not living?”) or no encoding task. The current task was indicated by the color, font and case of the presented item. There were three conditions: no-task lists (participants did not have to perform judgments with the presented items), single-task lists (all items were presented with the same task), and two-task lists (items were presented with either task). In the two-task lists, items were presented successively with the same task in trains of 2–6 items, with train length chosen randomly. The first two lists were two-task lists, and each list started with a different task. The next fourteen lists contained four no-task lists, six one-task lists (three with each task), and four two-task lists. List and task order were counterbalanced across sessions and participants.

Each word was drawn from a pool of 1638 words (available at memory.psych .upenn.edu/files/wordpools/PEERS wordpool.zip). Lists were constructed such that varying degrees of semantic relatedness occurred at both adjacent and distant serial positions. Semantic relatedness was determined using the Word Association Space (WAS) model described by Steyvers, Shiffrin, and Nelson (2004). WAS similarity values were used to group words into four similarity bins based on the similarity between word pairs (high similarity, cos *θ >* 0.7; medium–high similarity, 0.4 *<* cos *θ <* 0.7; medium–low similarity, 0.14 *<* cos *θ <* 0.4; low similarity, cos *θ <* 0.14). Two pairs of items from each of the four groups were arranged such that one pair occurred at adjacent serial positions and the other pair was separated by at least two other items.

For each list, there was a 1500 ms delay before the first word appeared on the screen. Each item was on the screen for 3000 ms, followed by jittered (i.e., variable) interstimulus interval of 800–1200 ms (uniform distribution). If the word was associated with a task, participants indicated their response via a keypress. After the last item in the list, there was a jittered delay of 1200–1400 ms, after which a tone sounded, a row of asterisks appeared, and the participant was given 75 seconds to attempt to recall aloud any of the items from the most recent list.

### Electrophysiological recordings and data processing

Netstation was used to record EEG from Geodesic Sensor Nets (Electrical Geodesics, Inc.) with 129 electrodes. The signal from all electrodes was digitized at 500 Hz by either the Net Amps 200 or 300 amplifier and referenced to Cz. Prior to any data processing, recordings were rereferenced to the average of all electrodes except those with high impedance or poor contact with the scalp. To eliminate electrical line noise, a first order 2 Hz stopband Butterworth notch filter was applied at 60 Hz.

We excluded any recalls that occurred within 1000 ms of the next recall to prevent overlap of the neural activity between these recalls. In both the neural data and the behavioral data, we excluded recalls from output positions 1-3, as such recalls may reflect recall from short-term memory in immediate free recall (Kahana, 1996), and such earlier immediate recalls may have shorter latencies (Murdock & Okada, 1970; Kahana, 2012). We calculated spectral power from 42 of the 129 electrodes, including electrodes in regions established in successful memory encoding (Long et al., 2014; Long & Kahana, 2017; Weidemann et al., 2009): bilateral anterior superior (corresponding to dorsolateral prefrontal cortex), bilateral anterior inferior (corresponding to inferior frontal cortex), and bilateral posterior inferior (corresponding to inferior temporal cortex). From these electrodes, we calculated spectral power for each event (defined in the next paragraph) by convolving its EEG time series with Morlet wavelets (wave number = 6) at each of 46 frequencies logarithmically spaced between 2 Hz and 100 Hz.

For each frequency and electrode, power was averaged across the entire encoding or recall interval. Then, the power values were z-scored across encoding and recall events separately for each session to remove the effects of these variables. Thus, each study or recall event had a corresponding vector of z-scored power values, concatenated across 42 electrodes at each of the 46 frequencies.

We computed spectral power for defined events of interest: We defined encoding events as the time window from 200 ms to 3000 ms relative to the onset of each item’s presentation, and recall events as the time period -1000 ms to -600 ms relative to the verbalization of an item. The time window for presentation events was motivated by the choice of Manning et al. (2011), where the 200 ms delay was meant to account for the time delay between when the word appears on the screen and the participant begins to process the word, but otherwise activity is considered for the entire duration the word is on the screen. For the time window of recall events, we evaluated context reinstatement while varying the onset and duration of the time window. We evaluated time windows beginning from -1000 ms to -500 ms relative to the participant’s recall vocalization, ranging in duration from 300 ms to 800 ms (both ranges were assessed in increments of 100 ms). This evaluation of time windows indicated that context reinstatement was strongest for the recall time window of -1000 to -600 ms relative to recall vocalization.

### Feature selection

We followed the approach of Manning et al. (2011) to determine patterns of neural activity that change gradually with each studied word. First, we applied principal components analysis (PCA) to the set power values across electrodes and frequency bands contributing to each study or recall event, as described above, using control lists only (no-task or single-task lists). We excluded from subsequent analyses those principal components that failed to explain a substantial proportion of the variance according to the Kaiser criterion (Kaiser, 1960). Next, we quantified the extent to which each of the included principal components was autocorrelated, and only used those principal components meeting the threshold for being substantially autocorrelated across each word presentation. We follow the terminology of Manning et al. (2011) and refer to such principal components as *feature vectors*. To determine which of the components were feature vectors, for each feature *x* within each list *i*, we computed the Pearson’s lag 1 autocorrelation coefficient (*r_i_*) and associated *P* value. We then combined the autocorrelation coefficients across lists into a summary autocorrelation measure *r̄*:

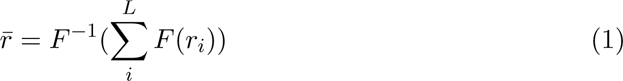

where *F* and *F^−^*^1^ are the Fisher z-prime transformation and Fisher inverse transformation, respectively. We also computed a summary measure for *P* across lists, *p̄*, by applying the inverse Normal transformation to the *P* values then summing across the transformed *P* values. We defined *p̄* as where the sum of the transformed *P* values fell on the the cumulative normal distribution function. Finally, we used *r̄* and *p̄* as inclusion criteria, and only included *x* as an autocorrelated feature vector if it satisfied *r̄ >* 0 and *p̄ <* 0.1.

The neural measure of temporal context was considered separately for each session. If there were not at least five autocorrelated feature vectors, the session was excluded from further neural and behavioral analysis. If the session did produce five autocorrelated feature vectors, we applied a PCA transformation matrix, determined from the control lists, to calculate temporal context vectors from two-task lists.

### Neural similarity

We defined the neural similarity between two feature vectors (in the participants’ data) or two temporal context vectors (in CMR) as the cosine of the angle between those two vectors (Manning et al., 2011). As confirmation of the approach for defining neural similarity during study, we calculated neural similarity as a function of study lag in one-task and two-task lists (Figure S1). For neural similarity during study of two-task lists, we only calculated similarity for pairs of items if both the preceding item and the following item were valid list positions (e.g. the first item on the list is not preceded by a list item, and thus was always excluded). We also excluded similarity values of preboundary items in events of 2 items, as the across-event similarity was included for the subsequent boundary item, and the within-event similarity was included for the preceding boundary item. We only calculated within-event similarity for pre-boundary and boundary items; otherwise, the within-event similarity measure would include many more items and may not be as comparable a comparison to across-event similarity. Finally, for preboundary items for which the across-event similarity was already included for the boundary item, we only included this value once as an across-event similarity value (i.e. we did not double-count these values).

For each recalled item, we calculated neural similarity only for lags that would be a valid recall transition. For instance, from serial position 1 it is not possible to make transitions of negative lags, and thus neural similarity for such items is not defined. We also did not calculate similarity between a recalled item and any of its study neighbors which were already recalled (this did not take into account the items excluded in output positions 1-3). Including neural similarity at study for a previously recalled item may be problematic, as neural similarity may reflect shared features from retrieval, not from study (cf. Folkerts et al., 2018). In the analysis of neural similarity for recalls from preboundary and boundary items in two-task lists, we only included the neural similarity values if the values were defined at both lag = +1 and lag = -1.

### Behavioral analyses

For comparable comparison between neural similarity and behavioral lag-CRPs during recall, we excluded the first three output positions from the lag-CRP analyses. Scripts used for calculating the lag-CRP while excluding these output positions is available at https://github.com/vucml/EMBAM.

### Statistical analyses

To compare between-participant conditions across participants with a large sample size, we used standard paired *t*-tests. All correlation analyses used robust regression, a regression measure less sensitive to potential outliers. Unlike standard Pearson’s regression, this analysis does not yield the same correlation and significance values if the dependent and independent measures are switched (i.e., the correlation of x and y is not the same as the correlation of y and x). In our analyses we defined the independent measure, plotted on the x axis, as the measure occurring earlier in time. Our regression analyses were motivated by CMR predictions, whereby a participant exhibiting a stronger impact of temporal disruption in neural reinstatement should also exhibit stronger temporal disruption during study, and stronger temporal disruption in recall behavior. Thus, for each of our regression analyses we had a hypothesized direction of the correlation, and we report one-tailed p-values.

## Acknowledgements

We thank Patrick Crutchley, Jonathan Miller, and Isaac Pedisich for assistance with programming the experiments and Adam Broitman, Elizabeth Crutchley, Kylie Hower, Joel Kuhn, and Logan O’Sullivan for help with data collection. We thank Michael Kahana for helpful discussions and sharing this dataset. This research was supported by NIH grant MH106266 to Lynn Lohnas and NIH grant MH055687 to Michael Kahana.

### Appendix Context Maintenance and Retrieval Model (CMR) simulations

Here we provide an overview of the CMR model, highlighting the components most relevant to interactions between temporal context and event segmentation. CMR stores representations of item features, their corresponding contexts, and the associations between items and context. When an item *i* is studied, this activates the item’s associated feature representation, **f***_i_*. CMR assumes a localist representation, such that item *i* is represented by a vector with 1 for those features corresponding to the item’s serial position and associated encoding task, and 0’s everywhere else. This feature representation is used to generate an input to update context, **c^IN^**_*i*_, using an association matrix that links item features (*F* ) to context states (*C*), *M^FC^*, with the simple product **c^IN^**_*i*_ = *M^F C^* **f***_i_*. This input to context is then used to update context: 

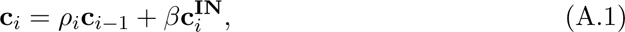

where *β* is a model parameter, and *ρ* is set so that *|***c***|* = 1 (a mathematical convenience). Larger values of *β* mean that the input to context will update context to a greater amount. When *β* is larger, *ρ* is smaller; as a result, the prior context is downweighed more. Note that in these equations, the index of each context and item feature is from the item *i*. Context is updated with each studied item, and thus changes slowly over time.

Critical to CMR’s ability to capture event segmentation, an event boundary updates temporal context beyond the updating from the studied item alone. Whenever there is a change in the task associated with an item, this causes CMR to present an additional ‘item’ to the model and update temporal context. However, these boundary items are not stored in memory and cannot be retrieved during the recall period. Nonetheless, they function to update context in a similar way to studied items, in that they update context according to Equation A.1. Whereas temporal context for a studied item is updated by setting *β* = *β^temp^_enc_*, temporal context for an event boundary item is updated with value *d*. This additional item thus disrupts the temporal context state, causing temporal context to drift even further from the current temporal context state. As a result, the temporal context between two items less similar when they are separated by an event boundary (Figure 3B).

Once CMR is presented with a list of items, the model next attempts to ‘recall’ items as a participant would. The model’s current state of context, reflecting the temporal history of studied items, is used to cue recall. Specifically, a feature strength is determined for each item, based on their relative weight in context: **f ^IN^_*r*_** = *M^CF^***c***_r_*, where elements of **f^IN^_*r*_** correspond to studied items. These feature strengths are then used as input to a noisy decision process that outputs a single recalled item, where items with larger strengths have a greater probability of being recalled (Usher & McClelland, 2001). As described in the next paragraph, because these feature strengths are determined from the current state of context, items with similar context states to the current context (i.e. items with shared temporal context or source context) are more likely to be recalled.

Once CMR recalls an item, this item is then presented to the model again, and updates context according to Equation A.1. Thus, a recalled item generates an input to context, and now this input includes the context from when the just-recalled item was originally studied. In addition, the rate at which context is updated, *β*, can vary between study and recall. Thus, the temporal context drift rates during study (*enc*oding) and *rec*all are termed and *β^temp^_enc_*and *β^temp^_rec_*, respectively. Once context is updated from the new item, this new state of context is used to recall another item. In this way, recall of an item *i* leads to reinstatement of the context of item *i*, and thus promotes recall of items with similar context states to *i*, including items with similar temporal contexts (i.e., items presented nearby on the list), as well as items with similar source contexts (i.e., items presented with the same task). This critical assumption of context updating during retrieval leads to CMR’s predictions of neural reinstatement (Figure 2C) and temporal contiguity in the behavioral lag-CRPs (Figure 2A).

Here we exclude details of CMR’s assumptions used to account for the increased recall of items studied at early list positions (primacy effect) and assumptions of pre-experimental semantic associations between words. These assumptions—while important to capturing recall dynamics—are less important in the current study where we average across serial positions and the lists contain mostly unrelated words (see Methods).

Instead of determining the parameter values that would best capture the present data, here we examined whether CMR could account for the data qualitatively based on best-fit parameters from a data set used previously (Polyn et al., 2009a). In this way, we were not fitting CMR to the data presented here, but rather using pre-existing parameter values and simulated data to predict the pattern of results for this data. To generate the CMR predictions, we presented the model with the same lists as participants viewed in that original study. Thus, the data used to generate CMR predictions included 45 participants with 631 control lists and 631 two-task lists each with list-length = 24.

**Figure S1:**
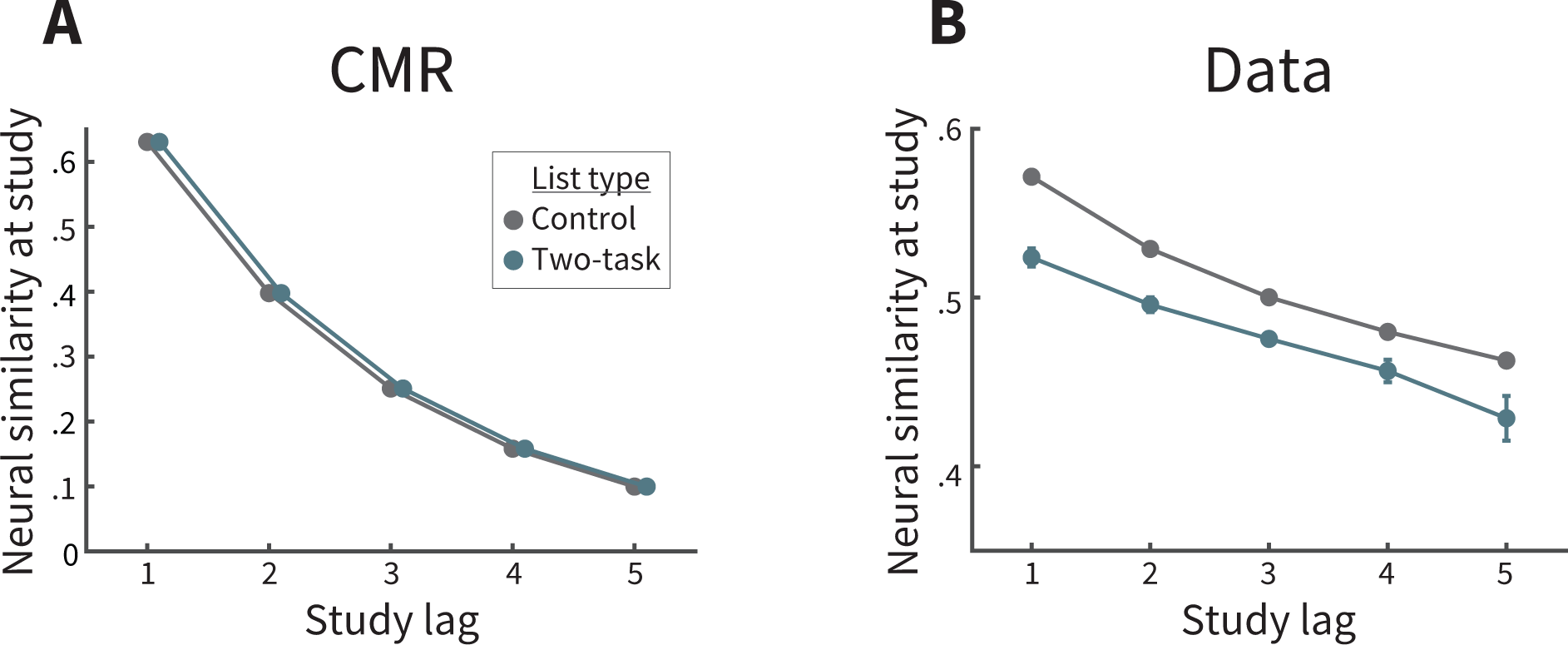
Neural similarity during study. **A.** CMR predicts that neural similarity n temporal context between two items should decrease as a function of lag. In two-task sts, items presented within the same event should have identical neural similarity values as n one-task lists. **B.** Participant data. As predicted by CMR (and as confirmation of our approach to calculate a neural measure of temporal context), neural similarity decreases as a unction of lag for both list types. That neural similarity is reduced in two-task lists reflects he choice of temporal context vectors from control lists and noisiness in the data. Error ars represent Loftus and Masson (1994) 95% confidence intervals.

## References

1. Aggleton, J. P., & Brown, M. W. (1999). Episodic memory, amnesia, and the hippocampal-anterior thalamic axis. Behavioral and Brain Sciences, 22, 425–489.

2. Baldassano, C., Chen, J., Zadbood, A., Pillow, J. W., Hasson, U., & Norman, K. A. (2017). Discovering event structure in continuous narrative perception and memory. Neuron, 95, 709–721. doi: 10.1016/j.neuron.2017.06.041

3. Blumenfeld, R. S., Parks, C. M., Yonelinas, A. P., & Ranganath, C. P. (2011). Putting the pieces together: The role of dorsolateral prefrontal cortex in relational memory encoding. Journal of cognitive neuroscience, 23, 257–265. doi: 10.1162/jocn.2010.21459

4. Chen, J., Leong, Y. C., Honey, C. J., Yong, C. H., Norman, K. A., & Hasson, U. (2017). Shared memories reveal shared structure in neural activity across individuals. Nature Neuroscience, 20, 115–125. doi: 10.1038/nn.4450

5. Clewett, D., & Davachi, L. (2017). The ebb and fl w of experience determines the temporal structure of memory. Current Opinion in Behavioral Sciences, 17, 186–193. doi: 10.1016/j.cobeha.2017.08.013

6. Clewett, D., DuBrow, S., & Davachi, L. (2019). Transcending time in the brain: How event memories are constructed from experience. Hippocampus. doi: 10.1002/hipo.23074

7. Clewett, D., Gasser, C., & Davachi, L. (2020). Pupil-linked arousal signals track the temporal organization of events in memory. Nature Communications, 11 (4007), 1–14. doi: 10.1038/s41467-020-17851-9

8. Davachi, L. (2004). The ensemble that plays together, stays together. Hippocampus, 14, 1–3. doi: 10.1002/hipo.20004

9. Davachi, L. (2006). Item, context and relational episodic encoding in humans. Current Opinion in Neurobiology, 16 (6), 693—700. doi: 10.1016/j.conb.2006.10.012

10. DuBrow, S., & Davachi, L. (2013). The infl of contextual boundaries on memory for the sequential order of events. Journal of Experimental Psychology: General, 142 (4), 1277–1286. doi: 10.1037/a0034024

11. DuBrow, S., & Davachi, L. (2014). Temporal memory is shaped by encoding stability and intervening item reactivation. The Journal of Neuroscience, 34 (42), 13998–14005. doi: 10.1523/JNEUROSCI.2535-14.2014

12. DuBrow, S., & Davachi, L. (2016). Temporal binding within and across events. Neurobiology of Learning and Memory, 134, 107–114. doi: 10.1016/j.nlm.2016.07.011

13. DuBrow, S., Rouhani, N., Niv, Y., & Norman, K. A. (2017). Does mental context drift or shift? Current Opinion in Behavioral Sciences, 17, 141–146. doi: 10.1016/j.cobeha.2017.08.003

14. Eichenbaum, H. (2004). Hippocampus: cognitive processes and neural representations that underlie declarative memory. Neuron, 44 (1), 109–120. doi: 10.1016/j.neuron.2004.08.028

15. Eichenbaum, H. (2014). Time cells in the hippocampus: a new dimension for mapping memories. Nature Reviews Neuroscience, 15 (11), 732–744. doi: 10.1038/nrn3827

16. Ezzyat, Y., & Davachi, L. (2011). What constitutes an episode in episodic memory? Psychological Science, 22 (2), 243–252. doi: 10.1177/0956797610393742

17. Ezzyat, Y., & Davachi, L. (2014). Similarity breeds proximity: Pattern similarity within and across contexts is related to later mnemonic judgments of temporal proximity. Neuron, 81 (5), 1179–1189. doi: 10.1016/j.neuron.2014.01.042

18. Faber, M., & Gennari, S. P. (2015). In search of lost time: Reconstructing the unfolding of events from memory. Cognition, 143, 193–202. doi: 10.1016/j.cognition.2015.06.014

19. Faber, M., & Gennari, S. P. (2017). Effects of event structure on prospective duration judgments. *Journal of Experimental Psychology: Learning*, Memory, and Cognition, 43 (8), 1203–1214. doi: 10.3758/BF03205466

20. Fernandez, G., Effern, A., Grunwald, T., Pezer, N., Lehnertz, K., Dumpelmann, M., . . . Elger, C. E. (1999). Real-time tracking of memory formation in the human rhinal cortex and hippocampus. Science, 285, 1582–1585. doi: 10.1126/science.285.5433.1582

21. Folkerts, S., Rutishauser, U., & Howard, M. W. (2018). Estimating scale-invariant future in continuous time. Journal of Neuroscience. doi: 10.1523/JNEUROSCI.2312-17.2018

22. Frank, N. T., Norman, K. A., Ranganath, C., Zacks, J. M., & Gershman, S. J. (2020). Structured event memory: A neuro-symbolic model of event cognition. Psychological Review, 127 (3), 327–361. doi: 10.1037/rev0000177

23. Goyal, A., Miller, J. F., Watrous, A. J., Lee, S. A., Coffey, T., Sperling, M. R., . . . Jacobs, J. (2018). Electrical stimulation in hippocampus and entorhinal cortex impairs spatial and temporal memory. Journal of Neuroscience, 19, 4471–4481. doi: 10.1523/JNEUROSCI.3049-17.2018

24. Hanslmayr, S., & Staudigl, T. (2014). How brain oscillations form memories– a processing based perspective on oscillatory subsequent memory effects. Neuroimage, 85, 648–655. doi: 10.1016/j.neuroimage.2013.05.121

25. Healey, M. K., & Kahana, M. J. (2014). Is memory search governed by universal principles or idiosyncratic strategies? Journal of Experimental Psychology: General, 143 (2), 575–596. doi: 10.1037/a0033715

26. Healey, M. K., Long, N. M., & Kahana, M. J. (2019). Contiguity in episodic memory. Psychonomic Bulletin & Review, 26 (3), 699–720. doi: 10.3758/s13423-018-1537-3

27. Heusser, A. C., Ezzyat, Y., Shiff I., & Davachi, L. (2018). Perceptual Boundaries Cause Mnemonic Trade-Offs Between Local Boundary Processing and Across-Trial Associative Binding., 44 (7), 1075–1090. doi: 10.1037/xlm0000503

28. Heusser, A. C., Poppel, D., Ezzyat, Y., & Davachi, L. (2016). Episodic sequence memory is supported by a theta–gamma phase code. Nature Neuroscience. doi: 10.1038/nn.4374

29. Howard, M. W., & Kahana, M. J. (2002). A distributed representation of temporal context. Journal of Mathematical Psychology, 46, 269–99. doi: 10.1006/jmps.2001.1388

30. Howard, M. W., Viskontas, I. V., Shankar, K. H., & Fried, I. (2012). Ensembles of human MTL neurons “jump back in time” in response to a repeated stimulus. Hippocampus, 22, 1833–1847. doi: 10.1002/hipo.22018

31. Hsieh, L.-T., Gruber, M., Jenkins, L., & Ranganath, C. (2014). Hippocampal activity patterns carry information about objects in temporal cortex. Neuron, 81 (5), 1165–1178. doi: 10.1016/j.neuron.2014.01.015

32. Jenkins, L. J., & Ranganath, C. (2010). Prefrontal and medial temporal lobe activity at encoding predicts temporal context memory. Journal of Neuroscience, 30 (46), 15558–15565.

33. Kahana, M. J. (1996). Associative retrieval processes in free recall. Memory & Cognition, 24 (1), 103–109. doi: 10.3758/BF03197276

34. Kahana, M. J. (2012). Foundations of human memory (1st ed.). New York, NY: Oxford University Press.

35. Kahana, M. J., Howard, M. W., & Polyn, S. M. (2008). Associative retrieval processes in episodic memory. In H. L. Roediger III (Ed.), Cognitive psychology of memory. Vol. 2 of Learning and memory: A comprehensive reference, 4 vols. (J. Byrne, Editor). Oxford: Elsevier.

36. Kaiser, H. F. (1960). The application of electronic computers to factor analysis. Educational and Psychological Measurement, 20 (1), 141–151. doi: 10.1177/001316446002000116

37. Kragel, J., Morton, N., & Polyn, S. (2015). Neural activity in the medial temporal lobe reveals the fi y of mental time travel. Journal of Neuroscience, 35 (7), 2914–26. doi: 10.1523/JNEUROSCI.3378-14.2015

38. Kurby, C. A., & Zacks, J. M. (2008). Segmentation in the perception and memory of events. Trends in Cognitive Sciences, 12 (2). doi: 10.1016/j.tics.2007.11.004

39. Loftus, G. R., & Masson, M. E. J. (1994). Using confidence intervals in within-subject designs. Psychonomic Bulletin & Review, 1, 476-490. doi: 10.3758/BF03210951

40. Lohnas, L. J., & Kahana, M. J. (2014). Compound cuing in free recall. *Journal of Experimental Psychology: Learning*, Memory and Cognition, 40 (1), 12–24. doi: 10.1037/a0033698

41. Long, N. M., Burke, J. F., & Kahana, M. J. (2014). Subsequent memory effect in intracranial and scalp EEG. NeuroImage, 84, 488–494. doi: 10.1016/j.neuroimage.2013.08.052

42. Long, N. M., & Kahana, M. J. (2017). Modulation of task demands suggests that semantic processing interferes with the formation of episodic associations. *Journal of Experimental Psychology: Learning*, Memory, and Cognition, 43 (2), 167–176. doi: 10.1037/xlm0000300

43. Long, N. M., Oztekin, I., & Badre, D. (2010). Seperable prefrontal cortex contributions to free recall. Journal of Neuroscience, 30 (33), 10967–10976. doi: 10.1523/JNEUROSCI.2611-10.2010

44. Lositsky, O., Chen, J., Toker, D., Honey, C. J., Poppenk, J. L., Hasson, U., & Norman, K. A. (2016). Neural Pattern Change During Encoding of a Narrative Predicts Retrospective Duration Estimates. eLife, 043075. doi: 10.1101/043075

45. MacDonald, C., Lepage, K., Eden, U., & Eichenbaum, H. (2011). Hippocampal “time cells” bridge the gap in memory for discontiguous events. Neuron, 71 (4), 737–749. doi: 10.1016/j.neuron.2011.07.012

46. Manning, J. R., Polyn, S. M., Baltuch, G., Litt, B., & Kahana, M. J. (2011). Oscillatory patterns in temporal lobe reveal context reinstatement during memory search. *Proceedings of the National Academy of Sciences*, USA, 108 (31), 12893–12897. doi: 10.1073/pnas.1015174108

47. Manns, J. R., Howard, M. W., & Eichenbaum, H. (2007). Gradual changes in hippocampal activity support remembering the order of events. Neuron, 56 (3), 530–40. doi: 10.1016/j.neuron.2007.08.017

48. McAndrews, M. P., & Milner, B. (1991). The frontal cortex and memory for temporal order. Neuropsychologia, 29 (9), 849–859.

49. Morton, N. W., Kahana, M. J., Rosenberg, E. A., Sperling, M. R., Sharan, A. D., & Polyn, S. M. (2013). Category-specific neural oscillations predict recall organization during memory search. Cerebral Cortex, 23 (10), 2407–2402. doi: 10.1093/cercor/bhs229

50. Murdock, B. B., & Okada, R. (1970). Interresponse times in singletrial free recall. Journal of Verbal Learning and Verbal Behavior, 86, 263-267. doi: 10.1037/h0029993

51. Norman, K. A., Polyn, S. M., Detre, G. J., & Haxby, J. V. (2006). Beyond mindreading: multi-voxel pattern analysis of fMRI data. Trends in Cognitive Sciences, 10 (9), 424–430. doi: 10.1016/j.tics.2006.07.005

52. Pacheco Estefan, D., Sanchez-Fibla, M., Duff, A., Principe, A., Rocamora, R., Zhang, H., . . . Verschure, P. F. M. J. (2019). Coordinated representational reinstatement in thehuman hippocampus and lateral temporal cortexduring episodic memory retrieval. Nature Communications, 10 (2255). doi: 10.1038/s41467-019-09569-0

53. Paller, K. A., & Wagner, A. D. (2002). Observing the transformation of experience into memory. Trends in Cognitive Sciences, 6 (2), 93–102. doi: 10.1016/S1364-6613(00)01845-3

54. Pettijohn, K. A., & Radvansky, G. A. (2018). Walking through doorways causes forgetting: recall. Memory, 26 (10). doi: 10.1080/09658211.2018.1489555

55. Pettijohn, K. A., Thompson, A. N., Tamplin, A. K., Krawietz, S. A., & Radvansky, G. A. (2016). Event boundaries and memory improvement. Cognition, 148, 136–144. doi: 10.1016/j.cognition.2015.12.013

56. Polyn, S. M., Norman, K. A., & Kahana, M. J. (2009a). A context maintenance and retrieval model of organizational processes in free recall. Psychological Review, 116, 129–56. doi: 10.1037/a0014420

57. Polyn, S. M., Norman, K. A., & Kahana, M. J. (2009b). Task context and organization in free recall. Neuropsychologia, 47, 2158–2163. doi: 10.1016/j.neuropsychologia.2009.02.013

58. Radvansky, G. A. (2012). Across the event horizon. Current Directions in Psychological Science, 21 (4), 269–272. doi: 10.1177/0963721412451274

59. Radvansky, G. A., & Copeland, D. E. (2006). Walking through doorways causes forgetting: situation models and experienced space. Memory & Cognition, 34 (5), 1150–1156. doi: 10.3758/BF03193261

60. Radvansky, G. A., Krawietz, S. A., & Tamplin, A. K. (2011). Walking through doorways causes forgetting: Further explorations. The Quarterly Journal of Experimental Psychology, 64 (8), 1632–1645. doi: 10.1080/17470218.2011.571267

61. Radvansky, G. A., & Zacks, J. M. (2014). Event cognition (1st ed.). New York, NY: Oxford University Press.

62. Radvansky, G. A., & Zacks, J. M. (2017). Event boundaries in memory and cognition. Current Opinion in Behavioral Sciences, 17, 133–140. doi: 10.1016/j.cobeha.2017.08.006

63. Richmond, L. L., & Zacks, J. M. (2017). Constructing experience: Event models from perception to action. Trends in Cognitive Sciences, 21 (12), 962–980. doi: 10.1016/j.tics.2017.08.005

64. Schapiro, A. C., Roger, T. T., Cordova, N. I., Turk-Browne, N. B., & Botvinick, M. M. (2013). Neural representations of events arise from temporal community structure. Nature Neuroscience, 16, 486—492. doi: 10.1038/nn.3331

65. Sederberg, P. B., Howard, M. W., & Kahana, M. J. (2008). A context-based theory of recency and contiguity in free recall. Psychological Review, 115 (4), 893–12. doi: 10.1037/a0013396

66. Sederberg, P. B., Miller, J. F., Howard, M. W., & Kahana, M. J. (2010). The temporal contiguity effect predicts episodic memory performance. Memory & Cognition, 38 (6), 689–699. doi: 10.3758/MC.38.6.689

67. Sederberg, P. B., Schulze-Bonhage, A., Madsen, J. R., Bromfield, E. B., Litt, B., Brandt, A., & Kahana, M. J. (2007). Gamma oscillations distinguish true from false memories. Psychological Science, 18 (11), 927–932. doi: 10.1111/j.1467-9280.2007.02003.x

68. Sols, I., DuBrow, S., Davachi, L., & Fuentemilla, L. (2017). Event boundaries trigger rapid memory reinstatement of the prior events to promote their representation in long-term memory. Current Biology, 27, 3499–3504. doi: 10.1016/j.cub.2017.09.057

69. Speer, N. K., & Zacks, J. M. (2005). Temporal changes as event boundaries: Processing and memory consequences of narrative time shifts. Journal of Memory and Language, 53 (1), 125–140. doi: 10.1016/j.jml.2005.02.009

70. Staresina, B. P., & Davachi, L. (2006). Differential encoding mechanisms for subsequent associative recognition and free recall. Journal of Neuroscience, 26 (36), 9162. doi: 10.1523/JNEUROSCI.2877-06.2006

71. Staresina, B. P., & Davachi, L. (2009). Mind the gap: binding experiences across space and time in the human hippocampus. Neuron, 63 (2), 267–276. doi: 10.1016/j.neuron.2009.06.024

72. Steyvers, M., Shiff R. M., & Nelson, D. L. (2004). Word association spaces for predicting semantic similarity effects in episodic memory. In A. F. Healy (Ed.), *Cognitive psychology and its applications: Festschrift in honor of Lyle Bourne, Walter Kintsch, and Thomas Landauer.* Washington, DC: American Psychological Association. doi: 10.1037/10895-018

73. Sugar, J., & Moser, M. (2019). Episodic memory: Neuronal codes for what, where, and when. Hippocampus, 29, 1190–1205. doi: 10.1002/hipo.23132

74. Swallow, K. M., Barch, D. M., Head, D., Maley, C. J., Holder, D., & Zacks, J. M. (2011). Changes in events alter how people remember recent information. Journal of Cognitive Neuroscience, 23 (5), 1052–1064. doi: doi.org/10.1162/jocn.2010.21524

75. Swallow, K. M., Zacks, J. M., & Abrams, R. A. (2009). Event boundaries in perception affect memory encoding and updating. Journal of Experimental Psychology: General, 138 (2), 236–257. doi: 10.1037/a0015631

76. Tsao, A., Sugar, J., Wang, C., Knierim, J. J., Moser, E. I., & Moser, M. (2018). Integrating time from experience in the lateral entorhinal cortex. Nature, 561, 57–62. doi: 10.1038/s41586-018-0459-6

77. Unsworth, N., Spillers, G. J., & Brewer, G. A. (2012). Dynamics of contextdependent recall: An examination of internal and external context change. Journal Of Memory And Language, 66, 1–16. doi: 10.1016/j.jml.2011.05.001

78. Usher, M., & McClelland, J. L. (2001). The time course of perceptual choice: The leaky, competing accumulator model. Psychological Review, 108 (3), 550–592. doi: 10.1037/0033-295X.108.3.550

79. Ward, G., Tan, L., & Grenfell-Essam, R. (2010). Examining the relationship between free recall and immediate serial recall: The effects of list length and output order. *Journal of Experimental Psychology: Learning*, Memory, and Cognition, 36 (5), 1207–1241. doi: 10.1037/a0020122

80. Weidemann, C. T., Mollison, M. V., & Kahana, M. J. (2009). Electrophysiological correlates of high-level perception during spatial navigation. Psychonomic Bulletin & Review, 16 (2), 313–319. doi: 10.3758/PBR.16.2.313

81. Zacks, J. M., Braver, T. S., Sheridan, M. A., Donaldson, D. I., Snyder, A. Z., Ollinger, J. M., . . . Raichle, M. E. (2001). Human brain activity time-locked to perceptual event boundaries. Nature, 4 (6), 651–655. doi: 10.1038/88486

82. Zacks, J. M., Speer, N. K., Swallow, K. M., Braver, T. S., & Reynolds, J. R. (2007). Event perception: a mind-brain perspective. Psychological Bulletin, 133, 273–293. doi: 10.1037/0033-2909.133.2.273

83. Zadbood, A., Chen, J., Leong, Y. C., Norman, K. A., & Hasson, U. (2017). How we transmit memories to other brains: constructing shared neural representations via communication. Cerebral cortex, 27, 4988–5000. doi: 10.1093/cercor/bhx202

84. Zwaan, R. A. (1996). Processing narrative time shifts. *Journal of Experimental Psychology: Learning*, Memory, and Cognition, 22 (5), 1196. doi: 10.1037/0278-7393.22.5.1196

